# Neuron-Microenvironments Associated with Specific Long-Range Connectivity Reveal Functional Modules in the Adult *Drosophila* Brain

**DOI:** 10.1101/2025.07.10.664244

**Authors:** Longxiao Yuan, Hanchuan Peng

**Affiliations:** New Cornerstone Science Laboratory, Institute for Brain and Intelligence, Fudan University, Shanghai, China; Shanghai Academy of Natural Sciences (SANS), Fudan University, Shanghai, China; School of Life Sciences, Fudan University, Shanghai, China

## Abstract

Advances in electron microscopy (EM) and computational neuroscience enable large-scale neuronal reconstruction, yet existing methods often overlook surrounding neurons’ influence. Using a whole-brain EM dataset of Drosophila, we analyzed 50 million synapses and derived microenvironmental (MicroEnv) features for over 130,000 neurons categorized by polarity. This refined 78 brain regions into 482 subregions, forming a high-resolution MicroEnv-atlas. MicroEnv features improved spatial coherence by 68% compared to single-neuron morphological features and correlated with long-range projections in the Mushroom Body. Integration with scRNA-seq data revealed associations with transmembrane transport and metabolism, suggesting roles in intercellular communication. Clustering based on distal projections identified submodules aligned with functional hierarchies, including feeding and multisensory processing. Our framework highlights the value of microenvironmental context in mapping brain organization.

## Introduction

Recent advances in high-resolution imaging have enabled the examination of brain architecture at single-neuron resolution. While traditional population-level labeling techniques have yielded valuable insights into neural circuitry, they often mask the diversity of individual projection patterns (Han et al., 2018; Zhou et al., 2023). In contrast, single-neuron analyses offer a powerful means to map connectivity with high specificity. Neural circuit function depends not only on the precise organization of long-range projections but also on dynamic interactions within local networks (Yonehara and Roska, 2016). Long-range axonal pathways are critical for behavioral modulation, and their disruption is frequently associated with neurological and psychiatric disorders (Dautan et al., 2024; Huang et al., 2023; Zhang et al., 2021). Accurate prediction of long-range connectivity is therefore crucial for uncovering circuit mechanisms and deepening our understanding of the neural basis of behavior and cognition.

Techniques such as MAP-seq, BAR-seq have aimed to link local microcircuitry with long-range axonal projections (Yonehara and Roska, 2016; Yuan et al., 2024), but their utility remains constrained by challenges like limited gene delivery efficiency and viral tropism (Chao et al., 2024; Li et al., 2018). These limitations underscore the need for refined predictive methodologies capable of resolving neuronal connectivity at single-cell resolution. Advances in computational methods have enabled large-scale reconstruction of neuronal morphology, facilitating the inference of long-range projections from structural features (Collman et al., 2015; Kasthuri et al., 2015). Data-driven approaches such as graph-based machine learning models encode complex morphologies into low-dimensional representations known as “morphological barcodes”, which have proven effective in distinguishing excitatory neuron subtypes in the visual cortex (Weis et al., 2025). In parallel, modeling strategies based on quantitative descriptions of axonal and dendritic probability density functions have shown promise in predicting projection patterns, particularly in hippocampal neurons (Gandolfi et al., 2023).

Intrinsic morphological features such as axonal arborization, spatial extent, and dendritic integration are strong predictors of projection patterns but often insufficient alone. To improve accuracy, recent approaches have combined molecular profiles with advanced computational models (Gao et al., 2023; Weis et al., 2025; Zhou et al., 2023). However, a critical factor remains underexplored: the influence of the neuron’s neighborhood. Beyond intrinsic traits, the local cellular milieu profoundly impacts neuronal survival, function, and connectivity (Li and Wu, 2025). Disruptions in this environment can alter cellular behavior and projection trajectories, highlighting the need for models integrating both intrinsic and extrinsic factors (Boedtkjer and Pedersen, 2020; Cheng et al., 2019). While the role of such a neighborhood, often called a microenvironment, is well established in cancer biology where it regulates tumor progression and therapy response, its impact on neuronal connectivity has received comparatively little attention. Incorporating microenvironmental context into predictive frameworks promises to enhance our understanding of brain circuitry and refine neuronal connectivity mapping.

To fill this gap, our group recently introduced the concept of the neuronal microenvironment in a study of the mouse brain. This framework integrates features derived from the surrounding local environment with intrinsic morphological descriptors to enhance predictive modeling (Liu, Y., et al., 2025). Our results demonstrate that incorporating microenvironmental information significantly improves the specificity of axonal projection predictions, particularly in the hippocampus, where projection patterns exhibit strong correlations with the structural characteristics of the local cellular context.

A goal of connectome research is to map all neuronal connections in the brain (Yonehara and Roska, 2016). Recently, researchers at Princeton University reached a major milestone by reconstructing the entire neuronal connectome of the adult female *Drosophila* brain using EM (Dorkenwald et al., 2024). The relatively small brain size allowed complete reconstruction of long-range projections. The dataset includes 139,273 neurons and approximately 50 million synapses, with hierarchical classification based on morphological features and spatial localization (Schlegel et al., 2024). This comprehensive whole-brain neuronal skeleton offers a unique opportunity to integrate microenvironmental context into the study of the Drosophila brain’s structural and functional architecture.

In this study, we computationally extracted microenvironmental features for all neurons and used these features to refine the subdivision of 78 anatomical brain regions of *Drosophila* based on microenvironmental similarity and thus generated a new brain atlas MicroEnv-atlas, which has 482 regions. We evaluated the utility of these features by measuring their spatial coherence across the brain and by assessing their correlation with long-range projection patterns. Integration with ScRNA-seq data further suggested that the microenvironmental features capture underlying biomolecular interaction networks. Finally, to investigate higher-order brain organization, we averaged projection pattern matrices within each region of MicroEnv-atlas, followed by clustering. Our analysis revealed functionally interconnected submodules.

## Result

### Unipolar Neurons Dominate the *Drosophila* Brain and Exhibit Balanced Pre/Post-Synaptic Counts

We considered the recently released electron microscopy based whole *Drosophila* brain dataset (Dorkenwald et al., 2024). This dataset offers more details than previous studies, e.g. additional information on the Lamina and Ocella **(Figure 1A)**. We analyzed this dataset and found that of the 139,273 neurons, 3.3% lack cell bodies due to an error during voxel-to-skeleton file conversion. Among the remaining neurons, 96.7% contain cell bodies. Unipolar neurons account for 92.7%, bipolar neurons for 6.9%, and multipolar neurons for 0.4% **(Figure 1B)**. Examples for each category are shown in **Figure 1D-F**.

**Figure 1.**
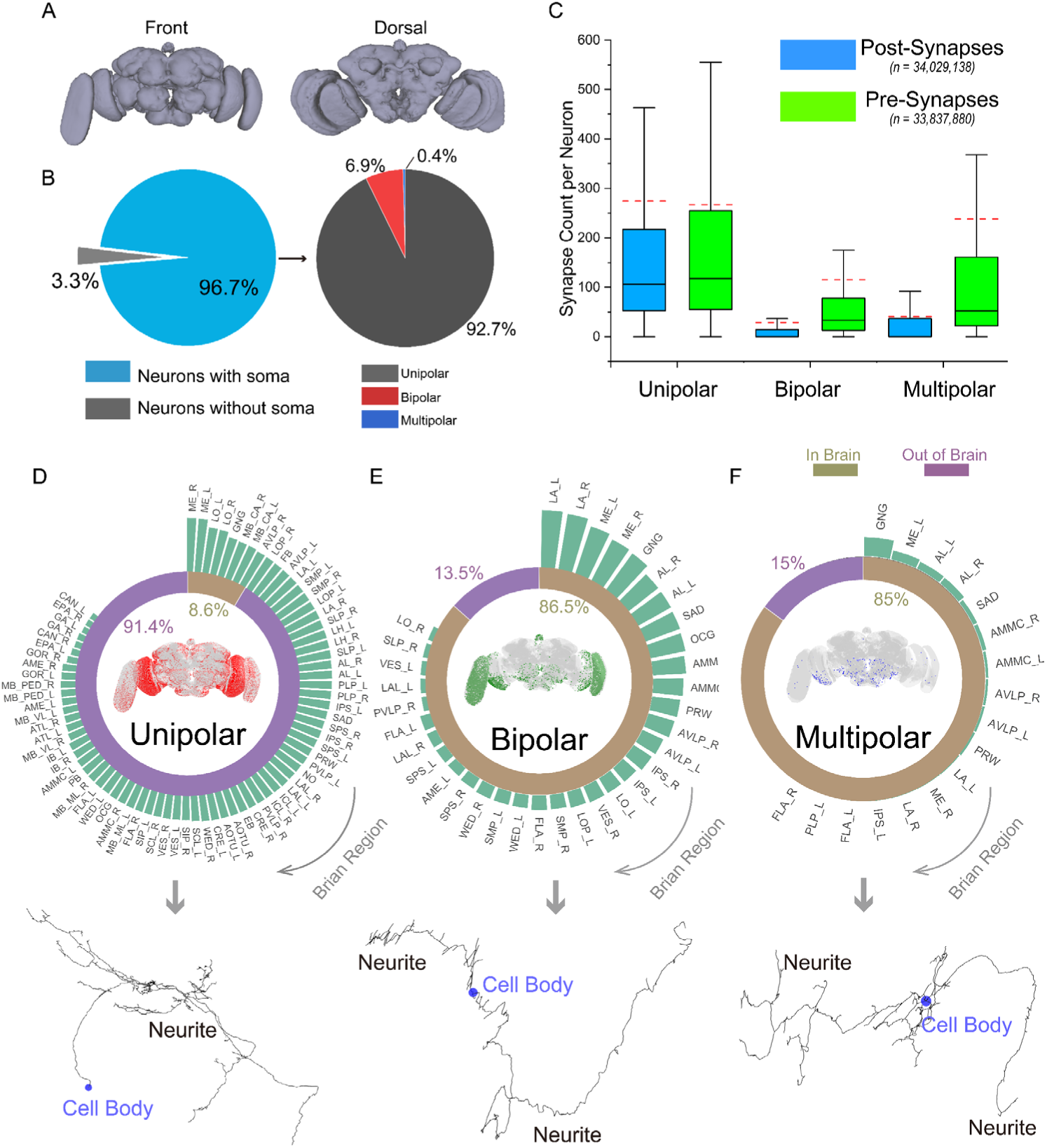
Overview of Drosophila whole-brain neurons. **A.** Drosophila whole-brain atlas shown in front and dorsal views. **B.** Proportional distribution of different neuron types across the entire brain. **C.** Box plots illustrating the distribution of input and output synapses across unipolar, bipolar, and multipolar neurons. Red dashed lines indicate the mean value for each group. **D-F.** Top panel: Radial bar chart showing the number of each neuron type across different regions of the Full Adult Fly Brain (FAFB) atlas. Circular pie chart depicting the proportion of neuron somas located inside (brown) versus outside (purple) the brain. Central panel: Anterior view of the Drosophila whole-brain outline, with scattered points representing neuron positions. Bottom panel: Representative skeletal structures of various neuron types, with blue circles marking cell bodies.

Unlike mammalian neurons, Drosophila neurons lack clearly demarcated dendrites and axons in the animal’s brain, and their pre- and post-synaptic sites are spatially interwoven with a relatively uniform degree of overlap across the brain **(Supplementary** Figure 3). To further investigate synaptic organization under this architecture, we quantified the distribution of pre-synaptic (output) and post-synaptic (input) membranes across different neuron types **(Figure 1C)**. The total number of synapses in unipolar neurons was more than 70 times greater than that of bipolar neurons and over 190 times greater than that of multipolar neurons. The average ratio of input to output synapses was 1.03, indicating a near-equal number of incoming and outgoing connections. This symmetry suggests that unipolar neurons exhibit a balanced distribution of synaptic input and output sites, potentially supporting stable signal transmission across the network. In contrast, bipolar and multipolar neurons exhibited average input-to-output synapse ratios of 0.25 and 0.17, respectively, with output synapses exceeding input synapses in both types. This asymmetry indicates a greater involvement in signal transmission rather than signal integration.

In addition to differences in total synapse count and input/output ratios, there are marked disparities in the spatial distribution of neurons. Unipolar neurons, the most abundant, are widely distributed, with the highest concentrations in ME, LO, and GNG. Bipolar neurons are concentrated in LA, ME, and GNG. Multipolar neurons, the least abundant, are primarily located in GNG. Further analysis of neuron cell body locations (*n = 134,689*) revealed that over 85% of these neurons are positioned outside the brain parenchyma (*n = 115,499*) **(Supplementary** Figure 4A). This proportion varies by neuron polarity: 91.4% of unipolar neurons are located outside the parenchyma, while bipolar and multipolar neurons have their cell bodies mostly within the parenchyma, at 86.5% and 85%, respectively **(Figure 1D-F & S4B-D)**.

### Neuronal microenvironment features reveal local spatial coherence in the *Drosophila* brain

As mentioned previously, only 134,689 neurons in the *Drosophila* brain skeleton files contained soma coordinates. We set a threshold of L/L_max_ = 0.15 to automatically correct errors at the soma connections. Of the neurons with soma, 130,485 were correctly annotated with brain regions. Given that the node distribution in the automatically generated neuron skeleton files was sparse and uneven, we resampled the skeletons using Vaa3D with a 1-μm sampling interval to address this imbalance and ensure that large neuronal skeletons could pass through subsequent feature extraction steps. This was followed by the extraction and calculation of 24 morphological features **(Figure 2B & Methods)**. Additionally, we computed 24-D microenvironment feature vectors based on the spatial distribution of neuron cell bodies **(Figure 2C & Methods)**. Subsequently, we examined how microenvironmental features varied across different neighborhood radii. We observed that once the radius exceeded the 30th percentile of distances to the sixth nearest neighbor across all neurons, the feature distributions exhibited regional consistency. Based on this finding, and to maintain consistency with previous studies (Liu, Y., et al., 2025), we adopted the 75th percentile of these distances as the neighborhood radius **(Supplementary** Figure 5).

**Figure 2.**
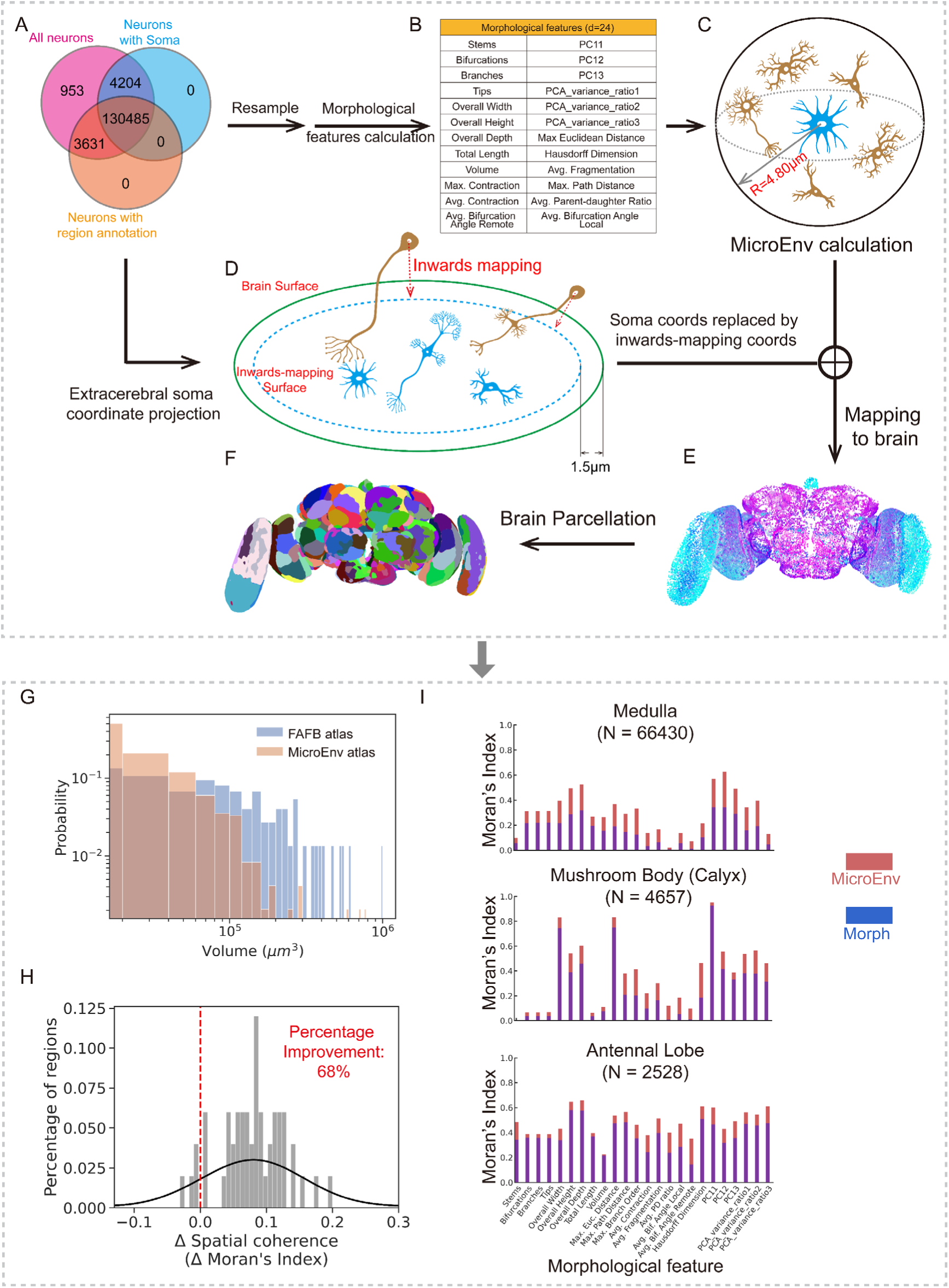
Construction and quality assessment of the MicroEnv-atlas. **A.** Venn diagram showing the intersection statistics of different neuron types in the Drosophila whole-brain dataset. **B.** Morphological features analyzed in this study (see **Methods** for details). **C.** Workflow for constructing the microenvironment. **D.** Method for inwards-mapping soma coordinates outside the brain onto the brain. **E.** Mapping microenvironment features onto the 3D whole-brain space. Microenvironments are color-coded in RGB based on the top three morphological features selected using the mRMR algorithm. **F.** MicroEnv-atlas constructed by segmenting the Drosophila brain based on microenvironment features. **G.** Histograms comparing the volume distribution of regions in the FAFB-atlas and the MicroEnv-atlas. Both the x-axis and y-axis are presented on an exponential scale for improved visualization. **H.** Distribution of spatial coherence changes (calibrated by Moran’s Index score) in FAFB-atlas after applying the microenvironment representation. The vertical red dashed line marks the threshold for non-improvement. **I.** Spatial coherence of morphological features across the brain.

Since over 85% of neuron cell bodies are located outside the brain parenchyma, we projected these cells into the brain volume to enable fine-grained segmentation based on the FAFB atlas **(Figure 2D, Supplementary** Figure 6A**, and Methods)**. To intuitively visualize the spatial distribution of microenvironments, we employed the top three microenvironmental features of each neuron **(Figure 2E)**. The top three features were identified with the minimum-Redundancy-Max-Relevance (mRMR) algorithm (Peng et al., 2005). The three features for the whole-brain feature map are the average of fragmentation (“Avg. Fragmentation”), corresponding variance ratios 1 (“pca_vr1”), and the number of main branches (“Stems”). Each inwards-mapping point or soma was treated as a node in an undirected graph, with an associated microenvironment feature vector. Nodes were classified based on spatial distance within the microenvironment, and regions with similar nodes were grouped into new brain regions. This process resulted in the fine segmentation of the *Drosophila* brain FAFB atlas into 482 subregions **(Figure 2F)**, forming the MicroEnv-atlas. The differences between subregions were significantly greater than the differences within regions **(Supplementary** Figure 6B).

We analyzed three matrices: volume, feature variability (“Feature STD”), and spatial autocorrelation (“Moran’s Index”). We found that volume was positively correlated with subregion count, while feature variability and Moran’s Index showed weak correlations with subregion count **(Supplementary** Figure 6C). Interestingly, subregions with larger feature variability and Moran’s Index values tended to have more subregions **(Supplementary** Figure 6D). This may be attributed to the inwards-mapping of cell bodies outside the brain into areas near the brain’s outline, leading to a denser distribution of coordinate points and higher consistency in connectivity, thus resulting in a higher number of subregions.

As expected, the subregions in the MicroEnv-atlas exhibited smaller volumes compared to those in the FAFB-atlas **(Figure 2G)**. Most MicroEnv-atlas subregions were confined within 10^5^ cubic micrometers, whereas larger subregions remained prevalent in the FAFB-atlas. For each FAFB region that was subdivided into subregions, the resulting subregion volumes exhibited moderate uniformity, as indicated by a Gini coefficient distribution that approximated a Gaussian distribution with a mean of 0.42 **(Supplementary** Figure 6E).

Compared to individual dendrites, microenvironments exhibited greater spatial coherence. Of the 78 FAFB regions, 68% showed increased Moran’s index values when microenvironmental context was incorporated **(Figure 2H)**. This was accompanied by reduced variance in intra-region features **(Supplementary** Figure 6F). Spatial coherence was especially pronounced in regions like Medulla, Mushroom Body and Antennal Lobe, where microenvironments demonstrated greater feature consistency compared to individual neurons **(Figure 2I)**.

### Microenvironment features in Mushroom Body correlate with neurons’ long-range projection

Predicting long-range neuronal projections and identifying key associated features is a central question in neuroscience, particularly for elucidating brain function and neural circuitry. In our previous work, we identified a strong correlation between neuronal microenvironmental (MicroEnv) features and long-range projections in the mouse brain, especially within the hippocampus (Liu, Y., et al, 2025). These findings motivated us to explore whether a similar relationship exists in the *Drosophila* brain.

To explore this, we calculated pairwise distances for the MicroEnv and projection features separately across various *Drosophila* brain regions and conducted Pearson correlation analyses. Overall, the correlations were relatively high in the Antennal Lobe, Mushroom Body, and Ocella. Notably, in both the Antennal Lobe and the Ocellar region, morphological characteristics showed stronger correlations with projection patterns. In contrast, within the Mushroom Body, microenvironmental attributes exhibited the highest correlation **(Figure 3A-B & Supplementary** Figure 7-8).

**Figure 3.**
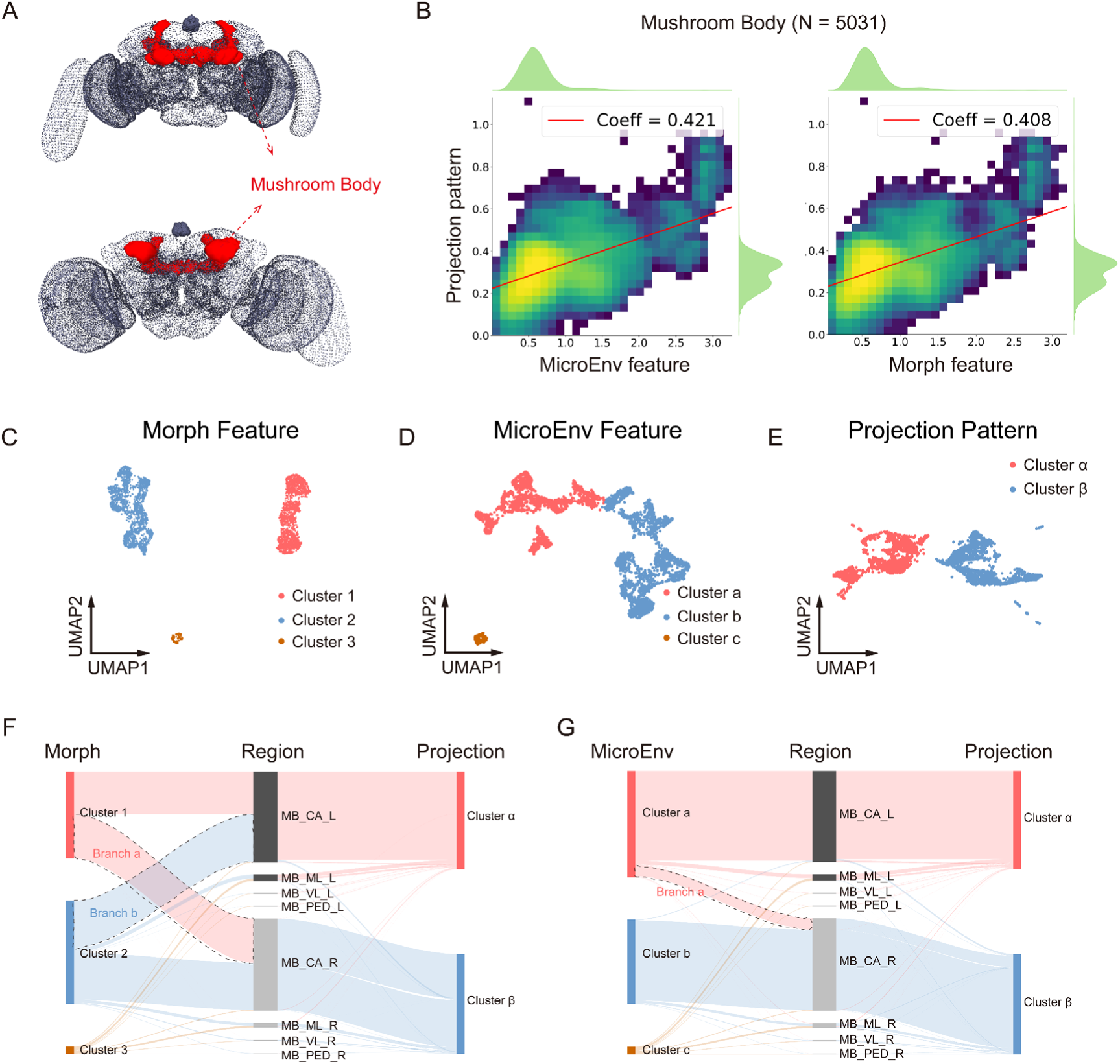
Correlation analysis between microenvironmental features and long-range projection features. **A.** Location of the Mushroom Body in the Drosophila brain, shown in front (top) and dorsal (bottom) views. **B.** Pearson correlation analysis of pairwise distances between neuronal MicroEnv features and axonal projection features in the Mushroom Body (left), and between neuronal Morphological features and axonal projection features (right). Color intensity from yellow to dark blue indicates decreasing neuron density. **C–E.** UMAP two-dimensional visualization of Morph, MicroEnv, and Projection features, with colors representing clusters identified via K-means clustering. **F.** Alluvial diagram illustrating the correspondence between Morph feature clusters (left), soma regions in the FAFB-atlas (middle), and axonal projection clusters (right). **G.** Alluvial diagram illustrating the correspondence between MicroEnv feature clusters (left), soma regions in the FAFB-atlas (middle), and axonal projection clusters (right), with cluster colors matching those used in the UMAP visualizations.

To better visualize the relationship between MicroEnv and Projection features in the Mushroom Body, we conducted dimensionality reduction on neuronal Morph, MicroEnv, and Projection features, followed by K-means clustering. Both Morph and MicroEnv features revealed three distinct clusters **(Figure 3C-D)**, while Projection features were grouped into two clusters **(Figure 3E)**. Using alluvial plots, we examined the flow of neurons across these clusters. Notably, Projection clusters α and β exhibited a strong correlation with the left and right Mushroom Body, respectively, with cluster α flowing toward the left hemisphere and cluster β toward the right hemisphere. This pattern was also reflected in the MicroEnv clusters, where clusters a and b primarily flowed toward the left and right Mushroom Body, respectively. However, cluster a showed a small branch (Branch-a) that deviated from the main flow and moved toward the right hemisphere **(Figure 3G)**. In stark contrast, Morph clusters 1 and 2 displayed no clear correspondence with the left and right hemispheres **(Figure 3F)**, with nearly half of the neurons from both clusters flowing to the opposite hemisphere. This difference is more apparent when visualized using the brain region-colored UMAP clustering plot **(Supplementary** Figure 9A-C).

Canonical correlation and linear regression analyses further supported these findings **(Supplementary** Figure 9D). Across the whole brain, the Mushroom Body was the only region in which microenvironmental features consistently showed stronger correlations with long-range projections than morphological features.

Since the Mushroom Body is linked to learning and memory, it is often considered the *Drosophila* equivalent of the hippocampus (Huang et al., 2024; Wong et al., 2023). The strong correlation between MicroEnv and projections in this region mirrors findings in mice, suggesting that neuronal microenvironments in learning-related brain regions may be evolutionarily conserved (Liu et al., 2024). This implies that different species may share similar regulatory networks for learning and memory.

### Transcriptomic Evidence for an Transmembrane transport – Intracellular metabolism Network Underlying Microenvironmental Regulation

To understand why microenvironmental features outperform morphological features in the *Drosophila* mushroom body, MicroEnv and Morph features were integrated with ScRNA-seq data. Correlation coefficients were calculated using the Cluster Mean Matching method (Deng et al., 2019), yielding *R_MicroEnv_* (the correlation between each gene and MicroEnv features) and *R_Single_*(the correlation between each gene and morphological features). The distribution of *R_MicroEnv_* exhibited a significantly higher gene density in the high-correlation range compared to *R_Single_*, indicating a marked difference between the two **(Figure 4A)**.

**Figure 4.**
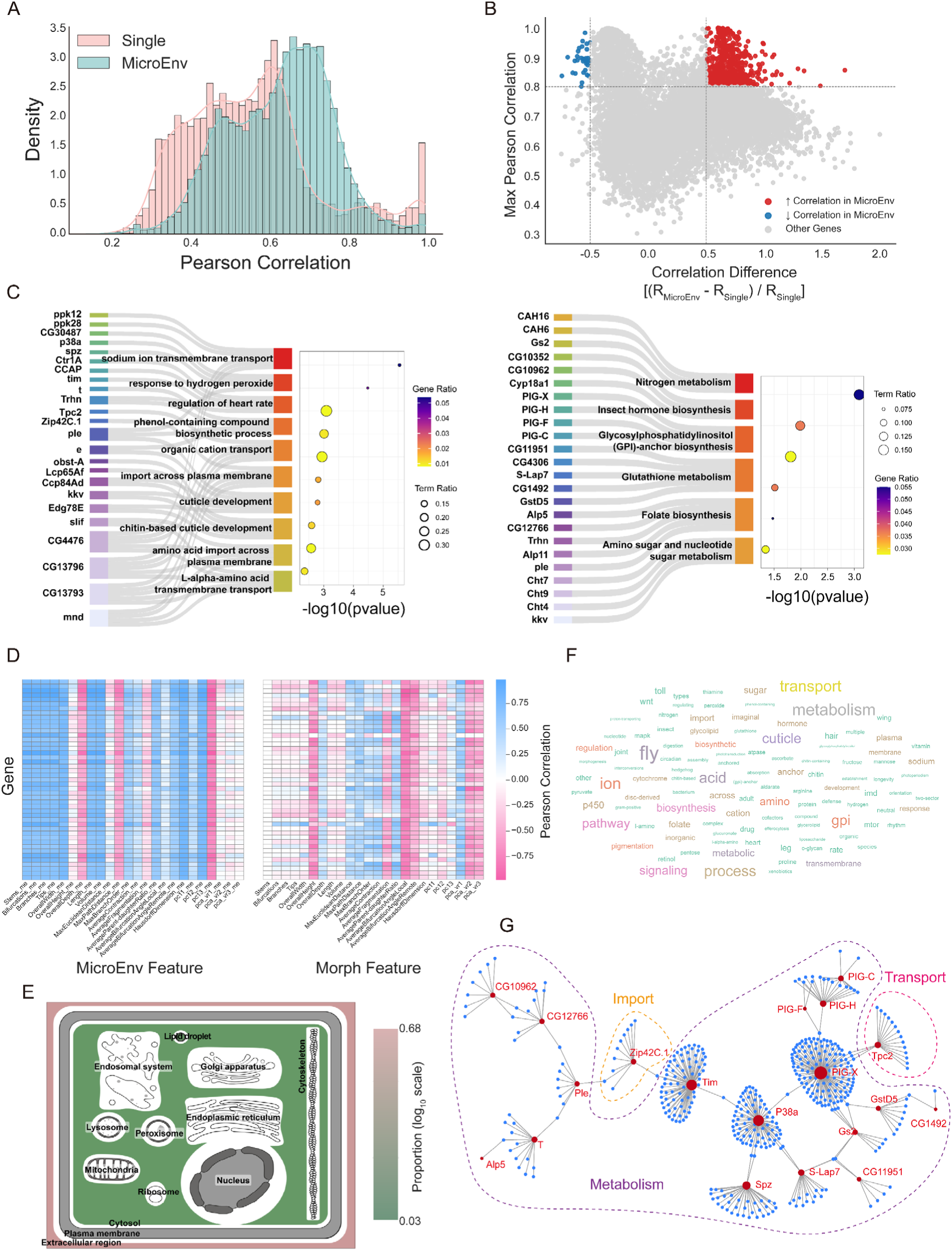
Correlation analysis between MicroEnv or morphological features and single-cell transcriptomic data. Let 𝑅_MicroEnv_ and 𝑅_single_ represent the Pearson correlation coefficients between each RNA transcript and the MicroEnv or morphological features, respectively. **A.** Distribution of 𝑅_MicroEnv_ and 𝑅_Single_ across all RNA transcripts. **B.** Volcano plot showing the difference (𝑅_MicroEnv_ – 𝑅_Single_) / 𝑅_Single_ on the x-axis and the higher of the two coefficients on the y-axis. Red dots indicate transcripts for which 𝑅_MicroEnv_ is significantly higher than 𝑅_Morph_, while blue dots indicate the opposite. Gray dots represent transcripts with no significant difference. **C.** GO (left panel) and KEGG (right panel) pathway enrichment analyses of genes with functional annotation among the red dots in **panel B**. Only pathways with p-values less than 0.05 are shown, with a maximum of 10 top-ranked pathways displayed. **D.** The Left Heatmap showing the correlation between each gene and individual MicroEnv features. The Right Heatmap showing the correlation between each gene and individual Morph features. **E.** Cellular component enrichment results of genes in the upregulated group. Genes are mapped to their respective subcellular localizations based on enrichment results, and each location is colored according to the proportion of enriched genes associated with it. **F.** Word cloud of the top-30 enriched pathway keywords from the GO/KEGG analyses. **G.** Protein – protein interaction (PPI) network of key genes. Red nodes represent top-enriched genes (genes that appear in **panel C**), while blue nodes represent other interacting proteins. Node size reflects the degree of connectivity.

To pinpoint which transcripts accounted for this divergence, a volcano plot was constructed. The x-axis represented the normalized difference, (*R_MicroEnv_* – *R_Single_*) / *R_Single_*, while the y-axis reflected the higher of the two correlation values. RNAs with x > 0.5 and y > 0.8 were classified as significantly upregulated in *R_MicroEnv_*, and those with x < −0.5 and y > 0.8 as significantly downregulated. This yielded 657 Gene in the upregulated group and 32 in the downregulated group **(Figure 4B)**. Functional enrichment analysis using GO and KEGG databases revealed a range of biological processes in upregulated group **(Figure 4C)**, while downregulated group lacked sufficient size to produce significant results. Additionally, correlation analysis of the top-enriched genes with individual MicroEnv and Morph features revealed strikingly distinct correlation patterns between the two feature sets, further highlighting the unique biological relevance of MicroEnv features **(Figure 4D)**.

The enriched pathways in the upregulated group were predominantly associated with material transport, metabolic processes, and biosynthesis, as revealed by GO and KEGG analyses. To further elucidate the spatial context of these biological functions, GO cellular component enrichment analysis was subsequently conducted. The top 10 enriched terms were mapped based on subcellular localization, revealing that most genes were associated with extracellular components **(Figure 4E & Supplementary** Figure 10). To better visualize the functional landscape, word clouds were generated for the top-30 GO and KEGG terms **(Figure 4F)**. Prominent keywords such as “transport”, “across”, “biosynthesis”, “transmembrane” and “metabolism” highlighted the involvement of these genes in intercellular exchange and cellular metabolism. Together, these findings support a conceptual model in which a self-regulating transport–metabolism loop facilitates microenvironmental communication.

To further explore this hypothesis, the top 10 enriched pathways were categorized into three functional classes: Import, Metabolism, and Transport (combined import/export) **(Supplementary Table 1).** A protein–protein interaction (PPI) network was then constructed using the top enriched genes **(Figure 4F)**. Network analysis revealed that many genes are functionally connected, either directly or indirectly, through protein interactions. Community labels based on pathway categories were applied to clusters within the network. Importantly, components from all four functional classes were present, further supporting the existence of an interconnected transmembrane transport – metabolism system underpinning microenvironmental communication in the mushroom body.

### The MicroEnv distribution of neurons in the *Drosophila* brain exhibits hemispheric asymmetry

A detailed segmentation of the Drosophila brain based on microenvironmental features led to the development of the MicroEnv-atlas, comprising 482 subregions. It was hypothesized that neuronal groups with similar projection patterns are functionally related or exhibit high internal consistency. To test this hypothesis, projection pattern vectors of neurons within each subregion were averaged, followed by cosine similarity-based clustering at multiple hierarchical levels. This analysis revealed two major modules, M1 and M2. Further subdivision of M2 resulted in submodules M2a and M2b, with M2b further divided into five finer groups: M2b1 through M2b5. Notably, these clustering outcomes closely corresponded to the previously established nine-category classification. **(Figure 5A)**.

**Figure 5.**
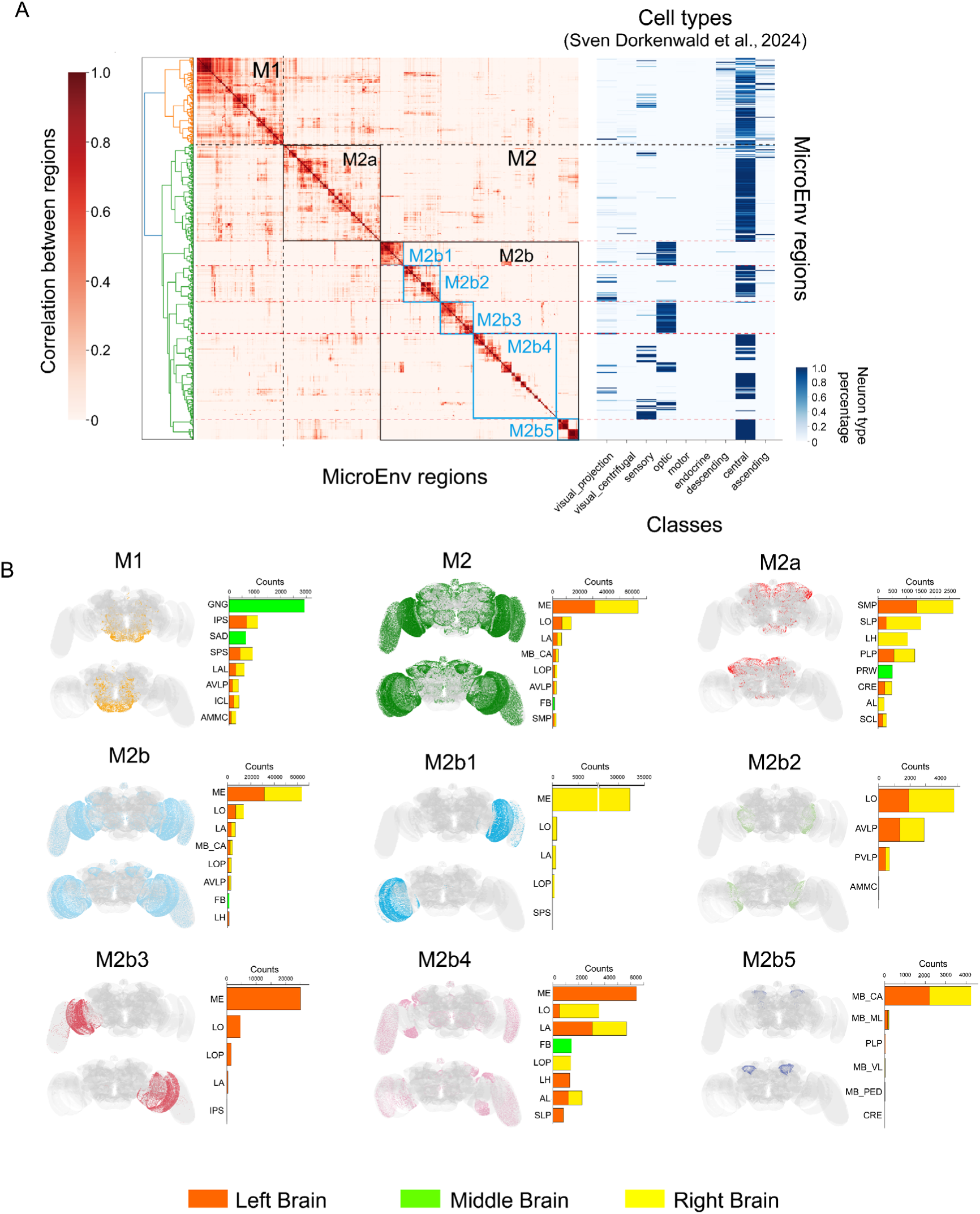
Projection pattern clustering reveals potential co-working modules. **A.** Left: Clustering of projection pattern similarities across MicroEnv-atlas. Right: Heatmap showing the distribution of neuron types (Flow hierarchy) across these regions. Modules (M1, M2) and submodules (M2a, M2b1, M2b2, M2b3, M2b4, M2b5) are outlined with solid squares, while dashed black lines indicate module boundaries. **B.** First column: Spatial distribution of neurons within each module in a 3D brain rendering (front and dorsal views). Second column: Neuron count statistics across brain regions, displaying up to the top eight regions. If fewer than eight regions are available, all will be shown. Left, middle, and right brain regions are color-coded in orange, green, and yellow, respectively.

Further analysis involved visualizing neurons from these submodules within a 3D brain map and statistically analyzing the distribution of neurons across various brain regions and hemispheres **(Figure 5B)**. The M1 module was predominantly localized to the GNG, IPS, and SAD regions, situated near the jaw area. In contrast, the M2 module, which comprised a larger population of neurons, was primarily enriched in visual regions. This spatial distribution aligns with the fruit fly’s highly developed visual system, which plays a critical role in survival and predator avoidance.

The M2a module was primarily located in the central brain, a region known for integrating and processing complex information (Kandimalla et al., 2023). The M2b module encompassed three major stages of visual processing: the lamina (LA), medulla (ME), and lobula (LO). Notably, the left and right hemispheres of the ME were mapped to M2b1 and M2b3, respectively, suggesting potential hemispheric differences in ME organization. The M2b2 module spanned the LO and central brain regions, specifically the anterior and posterior ventrolateral protocerebrum (AVLP and PVLP). This pattern supports the notion of consistent projection routes between LO and AVLP/PVLP, as LO is known to relay visual signals to the central brain via the lobula plate, thereby contributing to navigation-related behaviors (Stockl et al., 2020).

The M2b4 module included the optic lobes and parts of the central brain, encompassing the left and right hemispheres of LA and the left hemisphere of ME. These areas are responsible for early stages of visual processing. LA responds to changes in light intensity, while ME is involved in the initial formation of motion direction selectivity and the regulation of luminance gain (Gur et al., 2024; Pinto-Teixeira et al., 2018). The grouping of these regions into M2b4 suggests a direct communication pathway between the visual system and the central brain. This arrangement may facilitate rapid behavioral responses by reducing processing time and improving the efficiency of predator evasion. The final module, M2b5, was located in the mushroom body, a brain region that plays a central role in learning and memory in *Drosophila*.

Notably, the SLP region of the M2a module exhibited significantly fewer neurons in the left hemisphere compared to the right. A similar pattern was observed in the M2b4 module, where neurons were found exclusively in the left hemisphere of SLP and the right hemisphere of LH. Additionally, the right hemisphere of LO contained a greater number of neurons than the left. The ME region also demonstrated heterogeneous projection patterns, with neurons in the M2b5 module present in the left hemisphere of ME but absent from the right hemisphere. These observations suggest the presence of hemispheric asymmetry in neuronal projection patterns within the *Drosophila* brain. Such lateralized organization may contribute to the optimization of specific behaviors, including rapid escape responses and precise navigation, which are critical for survival and reproductive success. Alternatively, hemispheric heterogeneity might enhance the efficiency of neural information processing by minimizing redundancy across hemispheres. This organizational strategy could allow *Drosophila* to make more effective use of their relatively limited neuronal resources.

### Functional Interpretation of Brain Modules: Feeding and Multisensory Integration

Neurons in the *Drosophila* brain can be categorized into three types: Afferent (receiving external input), Intrinsic (internal integration), and Efferent (producing output) (Dorkenwald et al., 2024). This framework offers a useful model for interpreting the biological functions of the submodules we identified **(Figure 6A)**, where Afferent, Intrinsic, and Efferent neurons correspond to input, processing, and output components, respectively.

**Figure 6.**
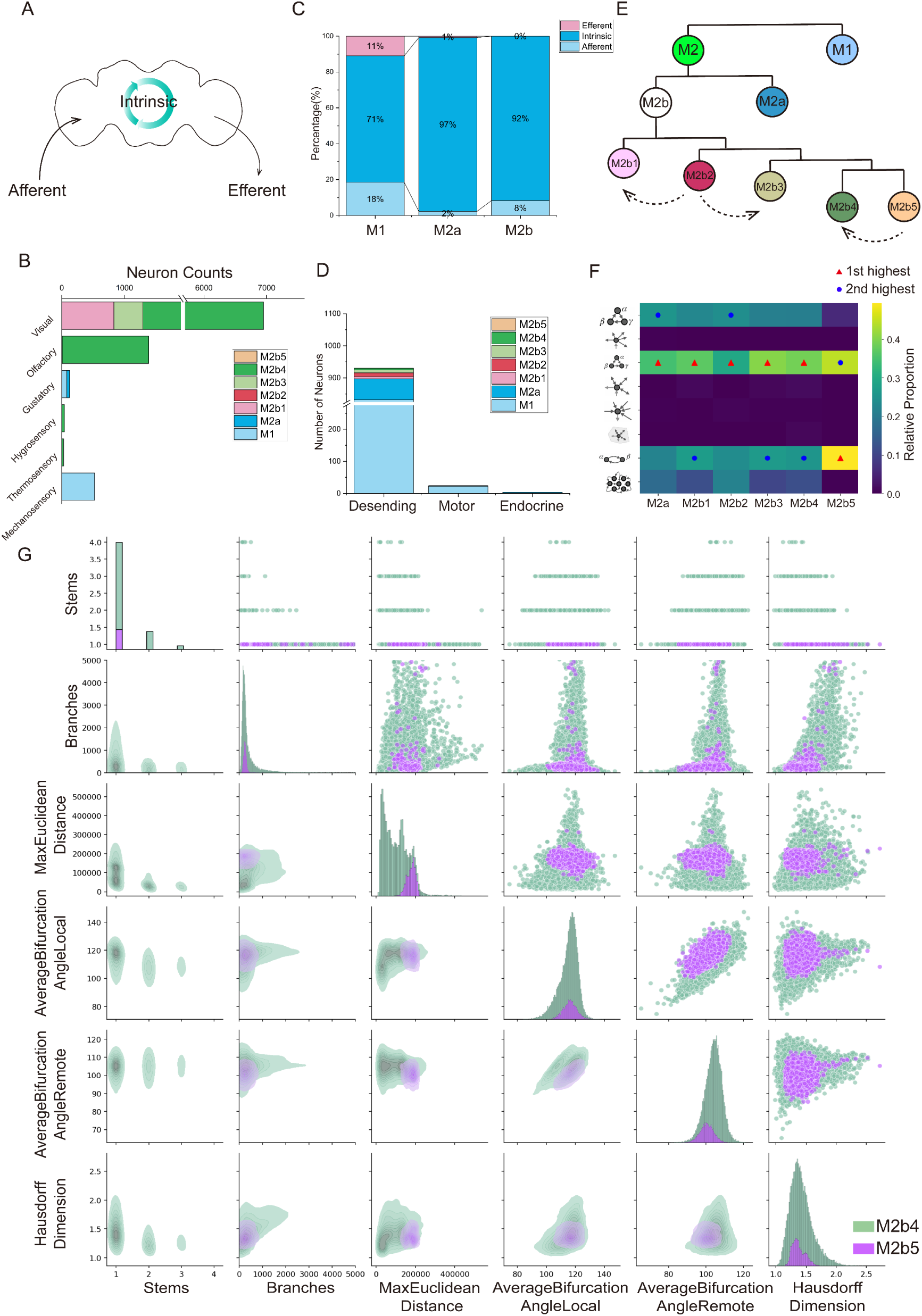
Biological functions associated with each module. **A.** Simplified model used for module functional identification. **B.** Proportion of efferent, intrinsic, and afferent neurons within modules M1, M2a, and M2b. **C.** Proportion of the 9 cell types in the sensory hierarchy assigned to each module. **D.** Proportion of the 3 cell types in the efferent hierarchy assigned to each module. **E.** Multilevel hierarchical clustering of functional modules, derived from the left panel of Figure 5A. Dashed arrows indicate potential collaborative relationships between modules, where the source module works for the target module. **F.** Composition of connectivity tags for neurons within modules M2a and M2b. Red triangles indicate the most abundant connectivity category, while blue circles indicate the second most abundant. The y-axis icons, from top to bottom, represent the following connectivity tags: 3-cycle participant, Broadcaster, Feedforward loop participant, Highly reciprocal neuron, Integrator, Neuropil-specific highly reciprocal neuron, Reciprocal connection participant, and Rich club. **G.** Pair plots of morphological feature distributions, with different colors representing the value distributions of neurons in each module.

We applied this classification to neurons in the M1, M2a, and M2b modules. M1 showed a relatively balanced composition: 18% Afferent, 71% Intrinsic, and 11% Efferent neurons. This suggests that M1 functions as a self-contained input-processing-output unit, likely carrying out specialized behaviors independently. M2a was dominated by Intrinsic neurons (97%), with only 2% Afferent and 1% Efferent neurons. This strongly indicates a role in internal computation, possibly supporting high-level sensory or cognitive processing. M2b also consisted mainly of Intrinsic neurons (92%) but had a slightly higher proportion of Afferent neurons (8%) and no Efferent neurons. Thus, M2b likely acts as an input-processing module **(Figure 6C)**.

To understand how each module contributes to behavior, we analyzed the input and output profiles of afferent and efferent neurons within each module. M1 primarily receives mechanosensory input, followed by gustatory signals **(Figure 6B)**, and projects to motor, descending, and endocrine targets **(Figure 6D)**. Its anatomical location near the jaw **(Figure 5B)** suggests that M1 integrates tactile and taste inputs to coordinate jaw movements and regulate feeding behavior.

Although M2a receives only gustatory input and includes some descending neurons, the predominance of intrinsic neurons suggests it functions as an information-processing module **(Figure 6B, 6D)**. In the hierarchical clustering tree, M2a is located near M2b **(Figure 6E)**, which implies that M2a may receive processed signals from M2b to refine behavioral decisions before generating output commands. Morphological analysis shows that M2a neurons have greater spatial extent and branching complexity, as indicated by higher Euclidean distances and Hausdorff dimensions, compared to M2b **(Supplementary** Figure 11). These features further support its role as a decision-making hub.

Within M2b, further submodule analysis revealed: M2b1 and M2b3 neurons are aligned along the optic pathway (ME), while M2b4 spans both the LA (visual input center) and AL (olfactory processing center) **(Figure 5B)**. These regions suggest M2b broadly integrates visual, olfactory, hygrosensory, and thermosensory inputs, with visual and olfactory inputs being dominant. M2b2 and M2b5 lacked direct sensory input, functioning instead as internal processing centers **(Figure 6B)**.

Specifically, M2b2 is located in the LO region **(Figure 5B)**, a critical relay for visual signals entering the central brain. Its close relationship with M2b1 and M2b3 **(Figure 6E)**, along with greater spatial complexity **(Supplementary** Figure 12-13), suggests that M2b2 specializes in deep visual information integration. M2b5 is located in the mushroom body **(Figure 5B)**, a region involved in memory, learning, and multisensory integration (Li et al., 2020; Zars, 2000). Given its proximity to the multisensory-integrating M2b4 **(Figure 6E)**, M2b5 likely supports higher-order processing. Morphologically, M2b5 neurons are more complex than those in M2b4, consistent with a role in refining integrated sensory signals **(Figure 6G)**.

To further validate these functional distinctions, we examined connectivity tags (Lin et al., 2024) assigned to neurons based on network topology. The submodules hypothesized to process external input (M2b1, M2b3, M2b4) shared similar tag patterns. In contrast, those involved in internal processing (M2a, M2b2, M2b5) showed distinct tag distributions **(Figure 6F)**. This pattern aligns with our previous interpretations and provides additional support for functional differentiation across modules.

## Discussion

In this study, we investigated the role of neuronal microenvironments in the structural and functional organization of the *Drosophila* brain. By leveraging the FlyWire dataset, we computationally derived microenvironmental features for all neurons and refined the subdivision of 78 anatomical brain regions based on microenvironmental similarity. Our findings confirmed that MicroEnv-atlas exhibit higher spatial coherence within regions. In correlation analyses with projection features, MicroEnv features showed advantages in Mushroom Body. Moreover, integration with ScRNA-seq data suggests that MicroEnv features may encapsulate information related to intercellular molecular communication. Furthermore, the clustering of projection patterns within microenvironment-based subdivisions revealed their potential for identifying functionally connected regions throughout the entire brain.

To enhance our understanding of the experimental findings, we analyzed the relationship between microenvironmental features and the long-range projection patterns of neurons. In the Mushroom Body, microenvironmental features exhibited stronger correlations with projection patterns than in other brain regions. This finding mirrors observations in the mouse hippocampus and is consistent with the known functional similarities between the Drosophila Mushroom Body and the mammalian hippocampus. Conversely, in regions outside the Mushroom Body, morphological features demonstrated a stronger correlation with projection characteristics. Notably, even the highest correlation observed in the Mushroom Body was only 0.421, which is significantly lower than the 0.752 reported for the mouse hippocampus (Liu et al., 2024). These differences highlight the need for further investigation into the underlying mechanisms.

Our results suggest that *Drosophila* neurons exhibit a widespread intermingling of pre-synapses and post-synapses throughout the brain, in stark contrast to the more segregated dendritic and axonal compartments found in mammalian neurons, such as those in mice (Liu et al., 2024). This structural difference may contribute to the generally weaker correlation between microenvironmental features and projection patterns compared to morphological features. Additionally, since pre-synapses and post-synapses annotations were obtained through automated computational identification, potential discrepancies from actual neuronal structures may introduce unavoidable biases in the correlation analysis.

Furthermore, research by Pogodalla et al. prompted us to consider another possibility. In *Drosophila*’s central nervous system, ensheathing glia encase the neuropil, limiting material exchange between the cell body and the neuropil. This creates distinct somatic and synaptic regions, which contrasts significantly with mammals (Pogodalla et al., 2021). Given the way we calculate microenvironmental features, it may be more accurate to refer to this as the soma region’s microenvironment. Perhaps it is the relatively independent developmental environments of the soma and synaptic regions under physiological conditions that contribute to this phenomenon.

Building on our observations of specific associations between microenvironmental features and long-range projection patterns, we further investigated whether these relationships could elucidate potential functional organization at the whole-brain level. To this end, we leveraged the previously constructed MicroEnv-atlas and integrated it with axonal projection characteristics to systematically identify functional modules. By clustering projection patterns based on cosine similarity, we identified several functional submodules within the MicroEnv-atlas, including M1, M2a, and M2b. These submodules exhibited a strong alignment with the hierarchical structure of neural information flow, underscoring the critical role of the neuronal microenvironment in shaping both information routing and the functional segmentation of the brain. This discovery advances our understanding of the functional topography of the *Drosophila* brain and suggests that neuronal connectivity patterns have been evolutionarily refined to support efficient information integration and behavioral control.

We propose that the M1 module is primarily associated with feeding behavior, potentially involving gustatory perception, and motor control. In contrast, the M2a and M2b modules are engaged in higher-level sensory integration and decision-making, likely coordinating cross-modal interactions such as vision, olfaction, and mechanosensation. This organization suggests that neuronal classification is not solely based on morphology but is also influenced by microenvironmental factors, leading to the formation of tightly coordinated functional communities. A similar design was observed in the study by Albert Lin et al., which found that neurons in the *Drosophila* brain tend to communicate within a “rich-club” structure. Specifically, interactions within hubs occur more frequently, while communication between different hubs is facilitated by only a few neurons serving as intermediaries. This finding suggests that the *Drosophila* brain operates in a modular fashion (Lin et al., 2024). Our results indicate that the presence of distinct modules reflects the brain’s optimization for various cognitive tasks, thereby enabling the execution of complex behaviors in an energy-efficient manner.

From an evolutionary perspective, small organisms like *Drosophila* must efficiently process information and make rapid decisions despite their limited energy resources (Ishimoto and Kamikouchi, 2021; Sareen et al., 2021; Stadele et al., 2019). Consequently, neural networks have evolved a modular topological structure that facilitates rapid processing within local networks while minimizing the energy costs associated with global communication. Notably, this microenvironment-based functional partitioning is not limited to the *Drosophila* brain but represents a general principle across various neural systems. For instance, similar functional module divisions are observed in the mammalian brain, such as the segregation of sensory, motor, and decision-making regions within the neocortex (Swanson et al., 2017; Vazquez-Rodriguez et al., 2019). This parallel suggests that neuronal microenvironments play a fundamental role in shaping the functionality of brain regions, representing a conserved organizational principle throughout evolution.

In summary, our findings underscore the crucial role of neuronal microenvironments in shaping both local neuronal properties and large-scale brain functional organization. This modular topology enables neural systems to efficiently process information and make adaptive behavioral decisions within resource constraints, presenting an evolutionarily optimized strategy for cognitive function. Furthermore, these insights not only advance our understanding of neural system organization but also have the potential to inspire novel approaches in artificial neural network design by integrating microenvironmental principles into computational models.

Despite these advances, our study is inherently limited by the available data. Currently, our analysis accounts solely for the neuronal microenvironment as defined by surrounding neurons, while critical components such as glial cells and mesenchymal cells remain uncharacterized due to data constraints. These non-neuronal elements play essential roles in neuronal function, synaptic plasticity, and metabolic support (Allen and Eroglu, 2017; Fernandes et al., 2024); their integration could provide a more comprehensive understanding of microenvironmental influences on neuronal behavior. We believe that incorporating a broader spectrum of microenvironmental information in future work could yield unprecedented insights into the fundamental principles underlying brain organization and function.

## Conclusion

This study demonstrates the value of microenvironmental features in refining the structural and functional organization of the *Drosophila* brain. By integrating spatial, morphological, and synaptic coordinate data, we constructed the MicroEnv-atlas, delineating 482 subregions and identifying functionally interconnected submodules. Notably, microenvironmental features outperformed morphological features in correlating with projection patterns within the Mushroom Body, possibly due to their capacity to capture intercellular molecular communication mediated by biomolecule exchange. In addition, submodules such as M1, M2a, and M2b correspond to information flow hierarchies, suggesting their involvement in feeding behavior and multi-sensory integration. Future work should incorporate additional contextual information surrounding neurons, including glial cells, vasculature, and metabolic factors, to further enrich the characterization of neuronal microenvironments. Extending this framework to other model organisms or human brain datasets may reveal conserved principles of neuronal organization. In summary, this study advances our understanding of brain architecture by highlighting the critical role of the microenvironment and provides a foundation for future research in systems neuroscience.

## Acknowledgment

This project was supported by several grants awarded to H.P. H.P. is a New Cornerstone Investigator and a SANS Senior Investigator. We would also like to thank Yufeng Liu, Feng Xiong, Zhixi Yun, and Xiaoqin Gu for their assistance in discussions.

## Author Contribution

H.P. conceptualized and designed the study and supervised the entire project. L.Y. performed the analyses, conducted the biological interpretation, and wrote the manuscript under the detailed guidance of H.P.. H.P. also contributed to the revision and provided critical guidance on the manuscript.

## Data and Materials Availability

The microenvironmental feature dataset has been archived at https://zenodo.org/records/15645086. The generated MicroEnv-atlas is also available in the same repository. Supporting data tables are provided as Supplementary Tables. The code used in the analysis can be accessed at https://github.com/ylx6266/MicroEnv/tree/main.

## Methods

### Nomenclature

The full names of the region abbreviations are: Accessory Medulla, AME; Lobula, LO; Nodulus, NO; Bulb, BU; Protocerebral Bridge, PB; Lateral Horn, LH; Lateral Accessory Lobe, LAL; Saddle, SAD; Cantle, CAN; Antennal Mechanosensory and Motor Center, AMMC; Inferior Clamp, ICL; Vest, VES; Inferior Bridge, IB; Antler, ATL; Crepine, CRE; Pedunculus of Mushroom Body, MB_PED; Vertical Lobe of Mushroom Body, MB_VL; Medial Lobe of Mushroom Body, MB_ML; Flange, FLA; Lobula Plate, LOP; Ellipsoid Body, EB; Antennal Lobe, AL; Medulla, ME; Fan-shaped Body, FB; Superior Lateral Protocerebrum, SLP; Superior Intermediate Protocerebrum, SIP; Superior Medial Protocerebrum, SMP; Anterior Ventrolateral Protocerebrum, AVLP; Posterior Ventrolateral Protocerebrum, PVLP; Wedge, WED; Posterior Lateral Protocerebrum, PLP; Anterior Optic Tubercle, AOTU; Gorget, GOR; Calyx of Mushroom Body, MB_CA; Superior Posterior Slope, SPS; Inferior Posterior Slope, IPS; Superior Clamp, SCL; Epaulette, EPA; Gnathal Ganglion, GNG; Prow, PRW; Gall, GA; Lamina, LA; Ocella, OCG.

### Fixing neuronal skeletons

The FlyWire team segmented the voxels of the entire brain of an adult female *Drosophila*, followed by manual corrections to each segmented neuronal voxel file (Dorkenwald et al., 2024; Schlegel et al., 2024). However, the automated conversion from voxel files to skeleton files introduced some unavoidable errors. Inaccuracies were observed in the reconstruction of soma nodes, characterized by radial patterns and abnormally large distances between the soma and the first connected node (Supplementary Figure 1A). To mitigate the impact of these inaccuracies, the maximum branch length (L) was quantified for each branch originating from the soma, with the longest branch defined as L_max_. The distance (D) from the soma to the first connected node was measured, with the distance for the longest branch labeled as D_max_. Analysis of the distribution differences in D and D_max_ across different L/L_max_ ratios revealed a significant shift when L/L_max_ exceeded 0.1. Consequently, a threshold of 0.15 for L/L_max_ was established to systematically remove erroneous branches (Supplementary Figure 1B).

### Annotating the polarity of neurons

We annotated neuronal polarity based on the corrected neuronal skeleton files. Neurons with a single branch were classified as unipolar, those with two branches as bipolar, and those with three or more branches as multipolar. During the evaluation of multipolar neurons, we observed that while most exhibited three distinct branches, some had short branches that did not extend beyond the radius of the soma (Supplementary Figure 2). Classifying these neurons as multipolar was clearly inappropriate. To ensure the accuracy of subsequent analyses, we manually reclassified these instances as bipolar using the SWC viewer tool, which we developed and is available for public use.

### Analysis of the Overlap in Pre-Synapse and Post-Synapse Distribution

We obtained the spatial coordinates of presynaptic and postsynaptic membranes for each neuron in the Drosophila brain from FlyWire dataset. To quantify axon-dendrite overlap, we defined a synapse as overlapped if at least three of its five nearest synapses belong to a different type. The degree of overlap is quantified as the proportion of overlapped synapses relative to the total number of synapses.

### Microenvironmental features extraction

We calculated 18 morphological features using the “global_neuron_feature” plugin of Vaa3D (Liang et al., 2023; Peng et al., 2014). These features include: “Stems”, “Bifurcations”, “Branches”, “Tips”, “Overall Width”, “Overall Height”, “Overall Depth”, “Total Length”, “Volume”, “Max. Euclidean Distance”, “Max. Path Distance”, “Max. Branch Order”, “Avg. Contraction”, “Avg. Fragmentation”, “Avg. Parent-daughter Ratio”, “Avg. Bifurcation Angle Local”, “Avg. Bifurcation Angle Remote”, and “Hausdorff Dimension”. Additionally, we performed principal component analysis (PCA) on all skeleton nodes to derive three principal component values (PC11, PC12, PC13) along with their corresponding variance ratios (pca_vr1, pca_vr2, pca_vr3), resulting in a 24-D morphological feature vector for each neuron. To ensure comparability across features, each feature was standardized using Z-score normalization, which scales the data to a common range, thereby eliminating differences in units and magnitudes and facilitating subsequent analyses.

In the FAFB space, a local spherical area with a radius of R = 4.80 µm (representing the 75th percentile of the distances to the sixth closest neighbor among all neurons) was defined around each target neuron (see Figure 2C). When more than six neurons were located within this sphere, the six nearest neurons to the target in the feature space were chosen; conversely, if fewer than six neurons were present, all neurons inside the sphere were included (refer to Eq. 1). The feature vectors of the neurons in the microenvironment were combined using spatial weighting, in which the weight assigned to each feature vector was based on the negative exponential of its relative distance from the target neuron, as explained in Eq. 2.

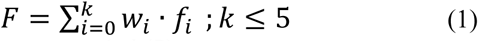

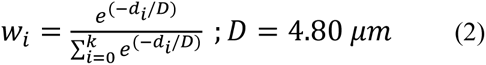

Where 𝑑 denotes the distance from the i-th neuron to the target neuron, 𝑓 represents the vector of morphological features, and 𝑘 indicates the total count of neighboring neurons associated with the target neuron.

### Brain parcellation

Since the majority of neuronal somas in the Drosophila brain are located outside the brain parenchyma, we ensured their inclusion in brain parcellation by projecting their coordinates onto the nearest brain surface, utilizing annotated brain region distribution data. These coordinates were subsequently shifted by 1.5 μm along the projection direction to integrate them into the brain structure. Neurons with soma depths ranging from 0 to 1.5 μm within the brain parenchyma account for less than 1.5% of the total, thereby minimizing interference with the shifted coordinates **(Supplementary** Figure 3A). We constructed an undirected graph based on the spatial distances within the microenvironment, adjusting these distances to account for regional shapes and calculating edge weights accordingly. The Leiden algorithm was employed to classify nodes, and communities were reassigned through majority voting, with parameters optimized using silhouette scores. Voxel communities were predicted using nearest-neighbor interpolation, smoothed through two rounds of 3D median filtering, and small subregions were removed and reassigned to ensure accurate and coherent parcellation.

### UMAP analysis

We reduced the 24-D microenvironmental features to 3 dimensions using UMAP (McInnes et al., 2018). The parameters of the UMAP model were configured as follows: the neighborhood size (n_neighbors) was set to 30 to balance the preservation of local and global structures; the minimum distance (min_dist) was set to 0.1 to control the density of points in the low-dimensional space; the dimensionality of the reduced space (n_components) was set to 2 to facilitate subsequent visualization and analysis; the distance metric (metric) was set to Euclidean distance (euclidean) to compute pairwise similarities in the high-dimensional space. Additionally, the learning rate (learning_rate) was set to 0.3, the spread of the embedding space (spread) was set to 1, and the initialization method (init) was set to the spectral method (spectral) to ensure the stability of the dimensionality reduction results. All other parameters were kept at their default settings.

### Pairwise-distance correlation analysis

We computed the pairwise distances for MicroEnv features, Morph features, and Projection features individually. Subsequently, we flattened the pairwise distance matrices and calculated the Pearson correlation coefficients between MicroEnv and Projection, as well as between Morph and Projection.

### Linear regression model analysis

We established linear regression models to analyze the relationships between MicroEnv features and Projection features, as well as between Morph features and Projection features across different brain regions. The dataset was randomly divided into training and testing sets in a 7:3 ratio. The R² score was employed to assess the model’s capability to explain the variance in the test dataset.

### Canonical correlation analysis

Canonical correlation analysis (CCA) was performed to examine the relationships between the MicroEnv features and Projection features, as well as between the Morphological features and Projection features, across various brain regions. The analysis was conducted with the hyperparameter set to n_components = 3 to capture the top three canonical components.

### Cosine similarity clustering

We calculated the average projection patterns across the 482 subregions defined by microenvironmental features. These average projection vectors were subsequently analyzed using cosine similarity and hierarchical clustering to identify connectivity patterns (Ward Jr, 1963). Additionally, we quantified the proportions of nine major neuronal classes (ascending, central, descending, endocrine, motor, optic, sensory, visual centrifugal, and visual projection) within each subregion, thereby offering insights into the functional composition of these regions.

### Cluster Mean Matching for Computing RSingle and RMicroEnv

To investigate the associations between transcriptomic features and both morphological and microenvironmental characteristics of cells, we developed a cross-modal correlation analysis framework based on Cluster Mean Matching. This approach quantifies the maximum linear correlation between each gene and either morphological features (R_Single_) or microenvironmental features (R_MicroEnv_) by aligning clustering structures across modalities.

Transcriptomic data (*GSE119629, GSE139889, GSE150642, GSE268425*), available in both TPM and count formats, were converted to count data using the Drosophila melanogaster gene annotation file (GTF format, BDGP6.46, Ensembl release 111). Morphological data, microenvironmental data, and transcriptomic data were first standardized. Specifically, all features were z-score normalized, followed by min-max scaling to the [0, 1] interval.

To construct a consistent topological structure across modalities, K-means clustering was performed independently on morphological, microenvironmental, and transcriptomic datasets using the same number of clusters (k = 10). For each modality, a cluster centroid matrix was computed, representing the mean value of each feature within each cluster.

For each gene, we quantified the similarity between its expression pattern and the distribution of morphological or microenvironmental features across clusters using the Pearson correlation coefficient. The procedure was as follows: 1. For each gene ***g***, its 10-dimensional expression vector across transcriptomic cluster centroids (**v*g***) was extracted; 2. Pearson correlation coefficients were computed between **v*_g_*** and the centroid vectors of all individual morphological features; 3. The maximum absolute correlation coefficient was retained as R_Single_ for gene ***g***; 4. The same procedure was applied using microenvironmental feature centroids to compute R_MicroEnv_.

### Enrichment Analysis and Word Cloud Visualization

To investigate the biological functions associated with genes that are upregulated in R_MicroEnv_, we conducted Gene Ontology (GO) enrichment analyses for biological processes (BP) and cellular components (CC), Kyoto Encyclopedia of Genes and Genomes (KEGG) pathway analysis, and keyword visualization (Ashburner et al., 2000; Aleksander et al., 2023; Kanehisa et al., 2025).

These genes were first preprocessed to convert gene symbols into standard identifiers compatible with enrichment tools. GO enrichment analysis was performed using the clusterProfiler package (Xu et al., 2024), with Entrez Gene IDs mapped via the *org.Dm.eg.db* annotation database. The analysis was restricted to biological process and cell component terms with a Benjamini –Hochberg adjusted p-value < 0.05. The top 10 significantly enriched BP GO terms were visualized using Sankey diagrams generated by GseaVis (Zhang et al., 2025). Similarly, KEGG pathway enrichment was conducted using the enrichKEGG function, specifying the Drosophila melanogaster organism code and retaining pathways with p < 0.05 and q < 0.2. Results were visualized using Sankey diagrams.

CC enrichment results were visualized with bar plots showing the top 10 enriched terms. In addition, a subcellular distribution map was generated based on the localization of each term and the proportion of associated genes. The initial subcellular localization sketch was derived from the *SubcellulaRVis* web tool (Watson et al., 2022).

To highlight the dominant biological themes, a word cloud was generated based on the top 30 GO and KEGG term descriptions. Terms were tokenized, cleaned, and filtered for length and stopwords. The ggwordcloud package was used to visualize word frequency using a circular layout, with word size and color proportional to term frequency.

### PPI Network Analysis

Top enriched genes in GO/KEGG were used with their official gene symbols. Protein–protein interaction analysis was performed using NetworkAnalyst-3.0 (https://networkanalyst.ca/NetworkAnalyst/) (Zhou et al., 2019), selecting the STRING Interactome database with a confidence score cutoff of 500.

---

**Supplementary Figure 1.**
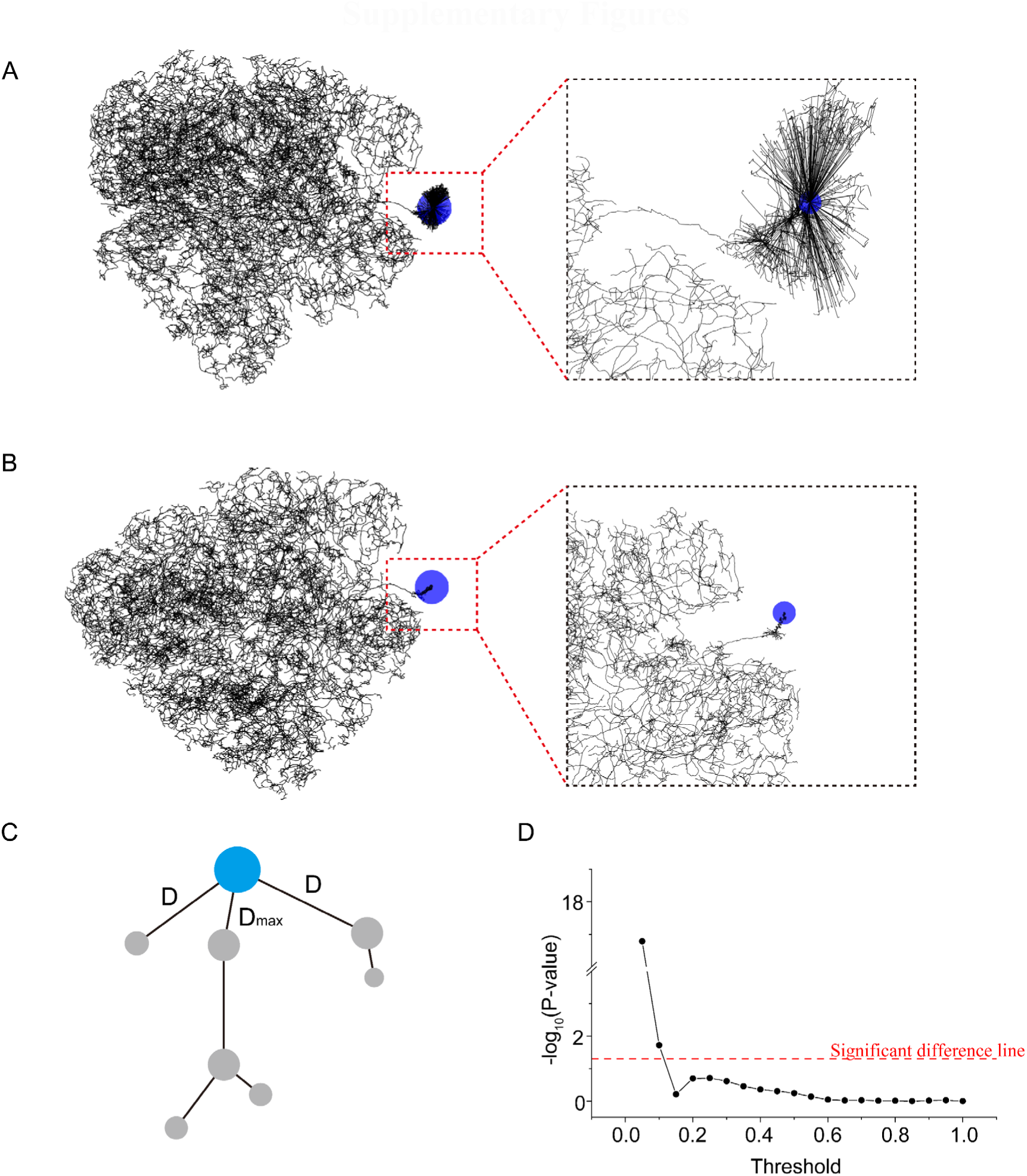
Correction of neuron skeletons. **A.** Example of a radial neuron at the soma connection. Left: Global view of the neuron; right: magnified view of the region within the red dashed box, with the soma highlighted in blue. **B.** Example of a corrected neuron skeleton after the fix. **C.** Diagram illustrating the simplified neuron skeleton: the distance from the soma to the first node is labeled as D, and the first segment of the longest branch is labeled as D_max_. **D.** Distribution differences of L and Lmax values under different threshold conditions (L_max_/L). The negative logarithm of p-values below the significant difference line indicates a significant difference (Mann-Whitney U Test) in the distribution of D and D_max_ values.

**Supplementary Figure 2.**
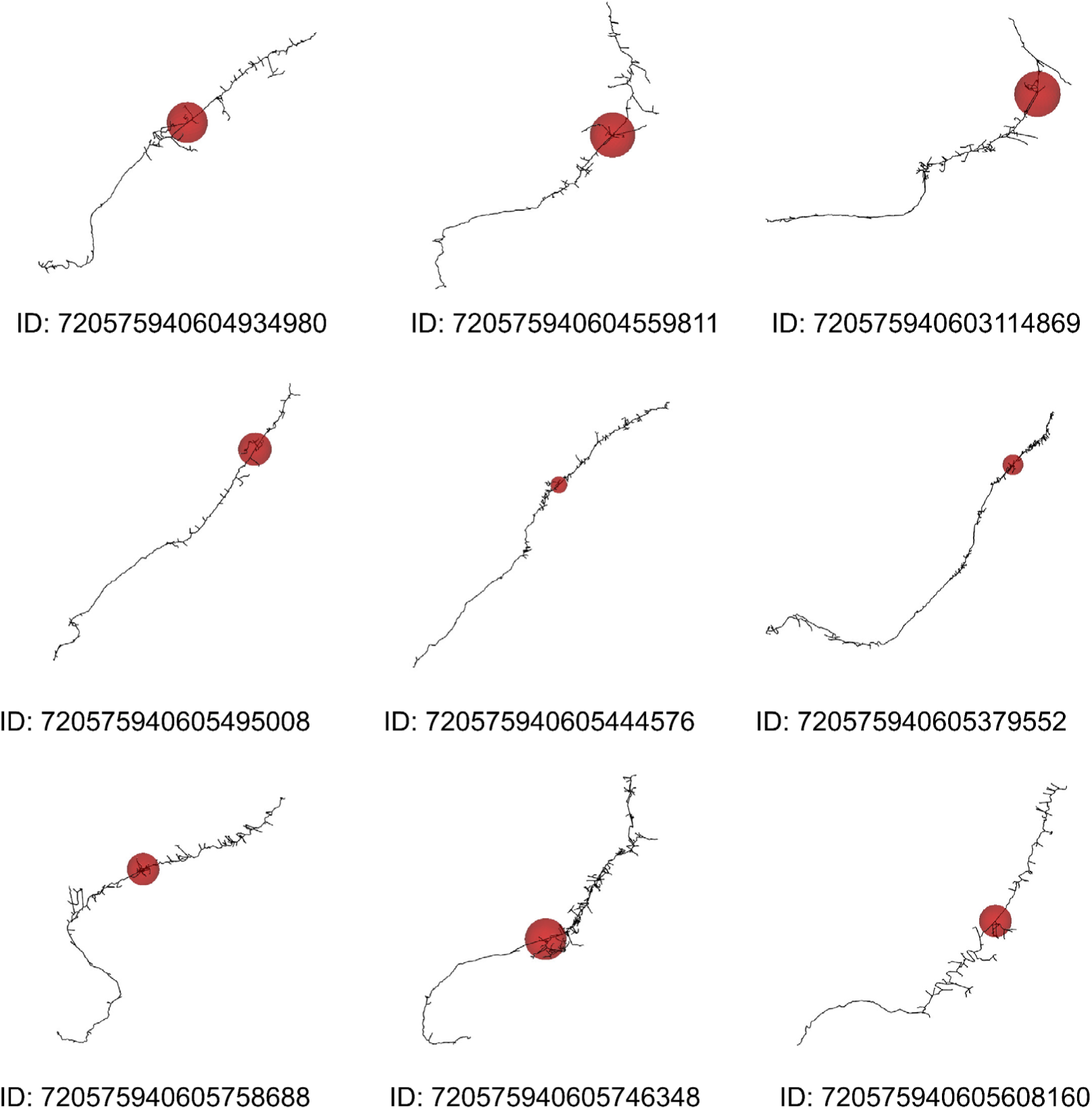
Example of bipolar neurons incorrectly classified as multipolar. Red spheres represent cell bodies, with the radius of each sphere reflecting their size.

**Supplementary Figure 3.**
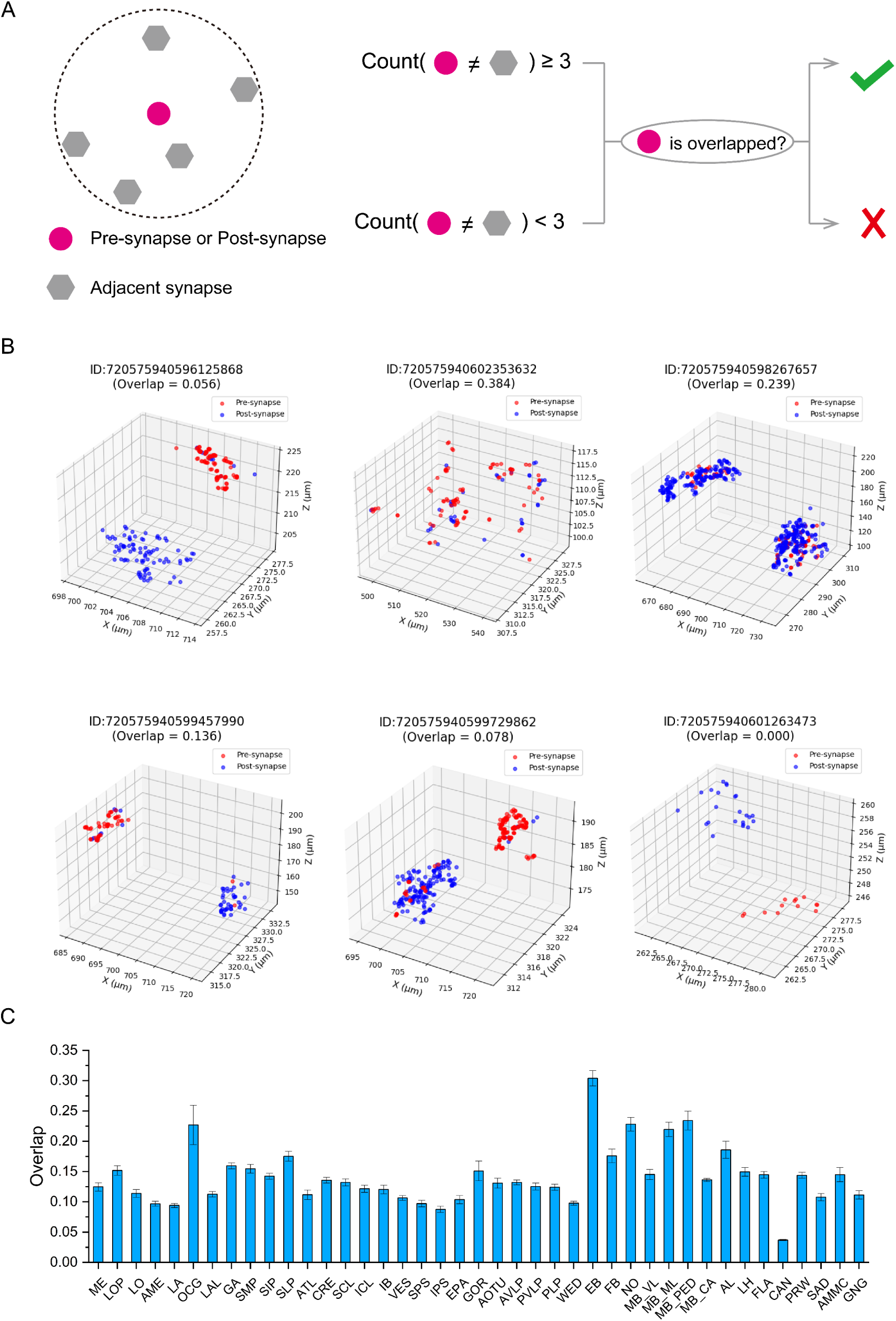
The Overlap in Pre-Synapse and Post-Synapse Distribution. **A.** Calculation method for determining the overlap between Pre-Synapse and Post-Synapse distributions. Red circles represent pre-synapses or post-synapses, while gray hexagons represent neighboring synapses. If three or more of the five neighboring synapses have a different type compared to the central synapse, the central synapse is considered overlapped. **B.** Examples showing the spatial distribution of pre-synapses and post-synapses in neurons, along with their spatial overlap. **C.** The degree of overlap between pre-synapses and post-synapses across different brain regions.

**Supplementary Figure 4.**
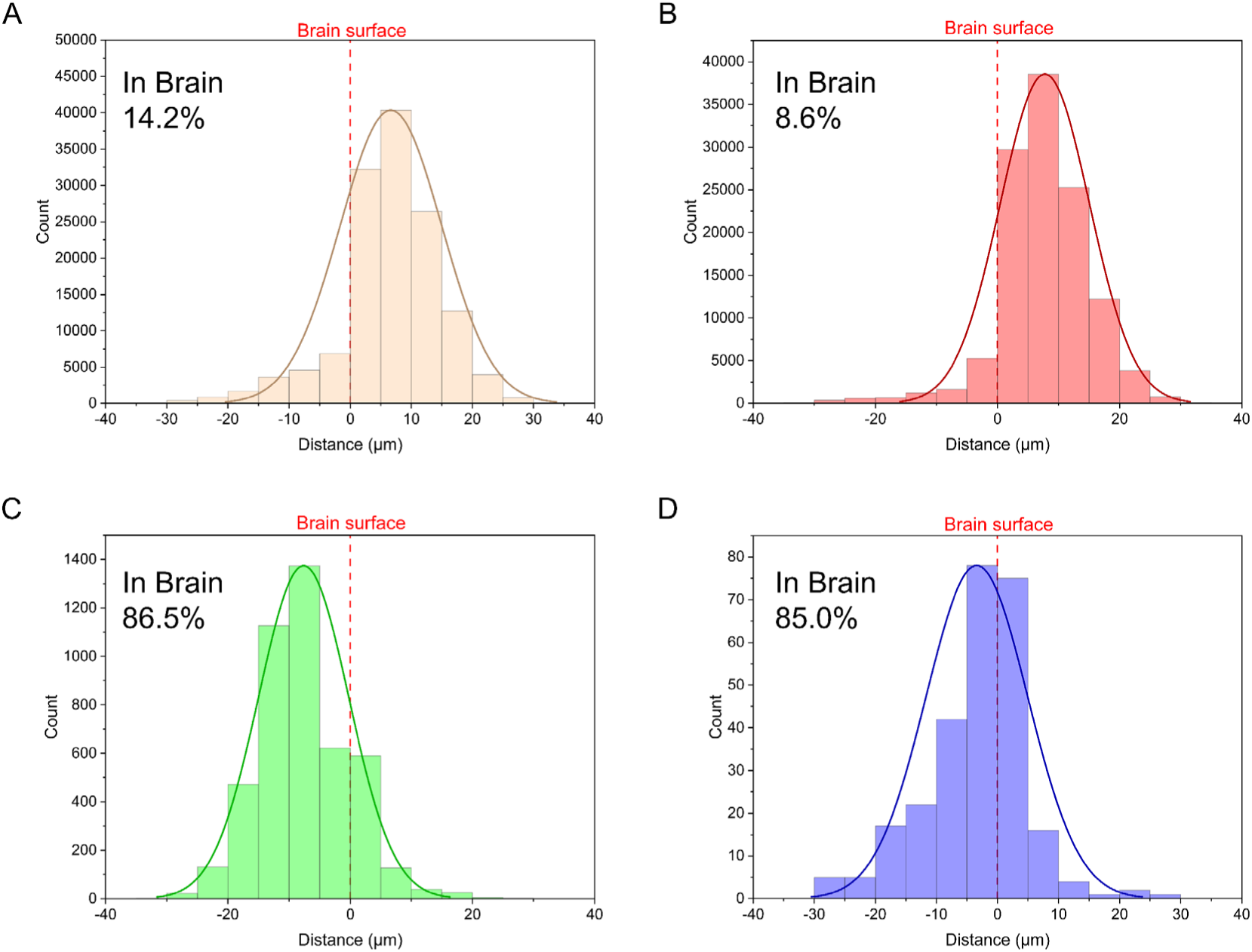
Soma-to-brain surface distance distribution for whole-brain and specific neuron types. **A-D.** Distance distributions for all neurons (A), unipolar neurons (B), bipolar neurons (C), and multipolar neurons (D). Positive values indicate soma locations outside the brain, while negative values indicate soma locations within the brain.

**Supplementary Figure 5.**
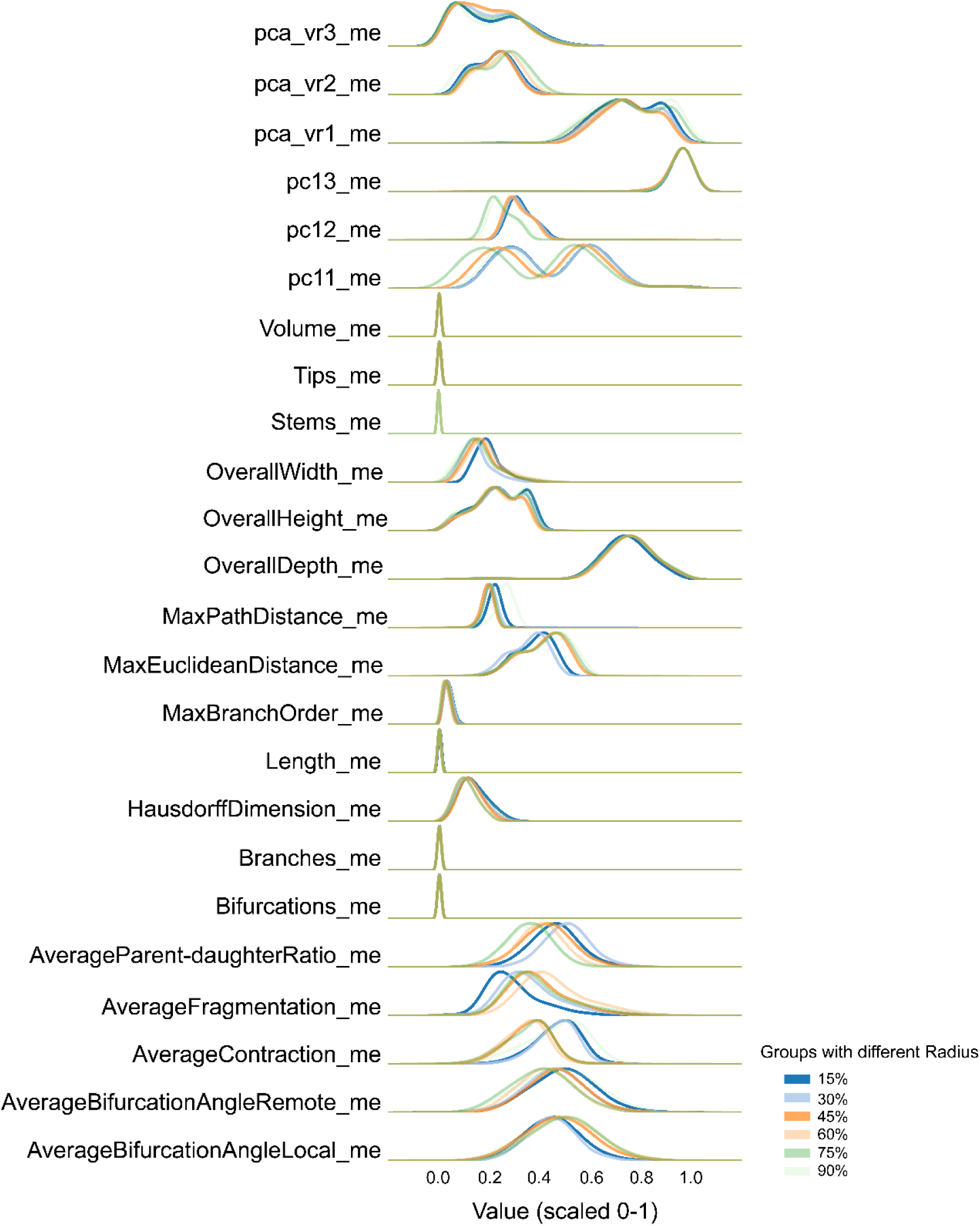
Distribution of Microenvironment Feature Values Under Different Radius Conditions. Value distributions (scaled from 0 to 1) of microenvironment features computed using different radius thresholds, defined as the 15%, 30%, 45%, 60%, 75%, and 90% quantiles of the distance to the sixth closest neighbor among all neurons.

**Supplementary Figure 6.**
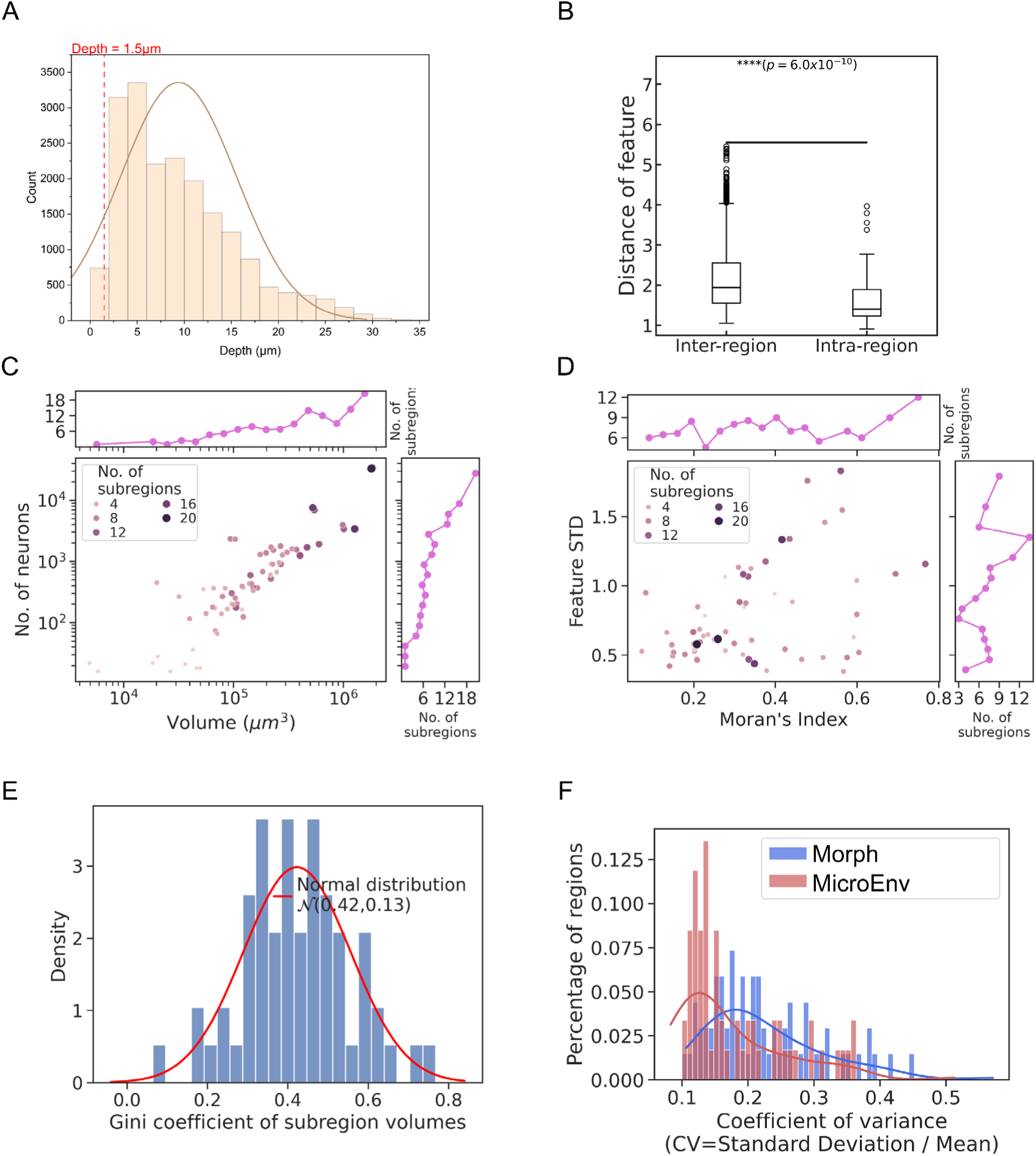
Microenvironment construction and brain parcellation quality assessment. **A.** Distribution of the number of neuron somas at different depths within the brain. **B.** Box plot showing inter-region and intra-region feature distances after brain segmentation. **C-D.** Relationship between the number of subregions and four metrics for each FAFB region. Left: Scatter plot of region volume (“Volume”) and number of neurons (“No. of neurons”) versus the number of subregions. Both features are plotted on a logarithmic scale. Right: Scatter plot of the standard deviation of microenvironment features (“Feature STD”) and spatial autocorrelation (“Moran’s Index”) versus the number of subregions. **E.** Histogram showing the Gini coefficients of subregion volumes within parcellated FAFB regions. A normal density function is fitted (μ=0.42, σ=0.13). **F.** Coefficient of variation distributions for morphology and microenvironments.

**Supplementary Figure 7.**
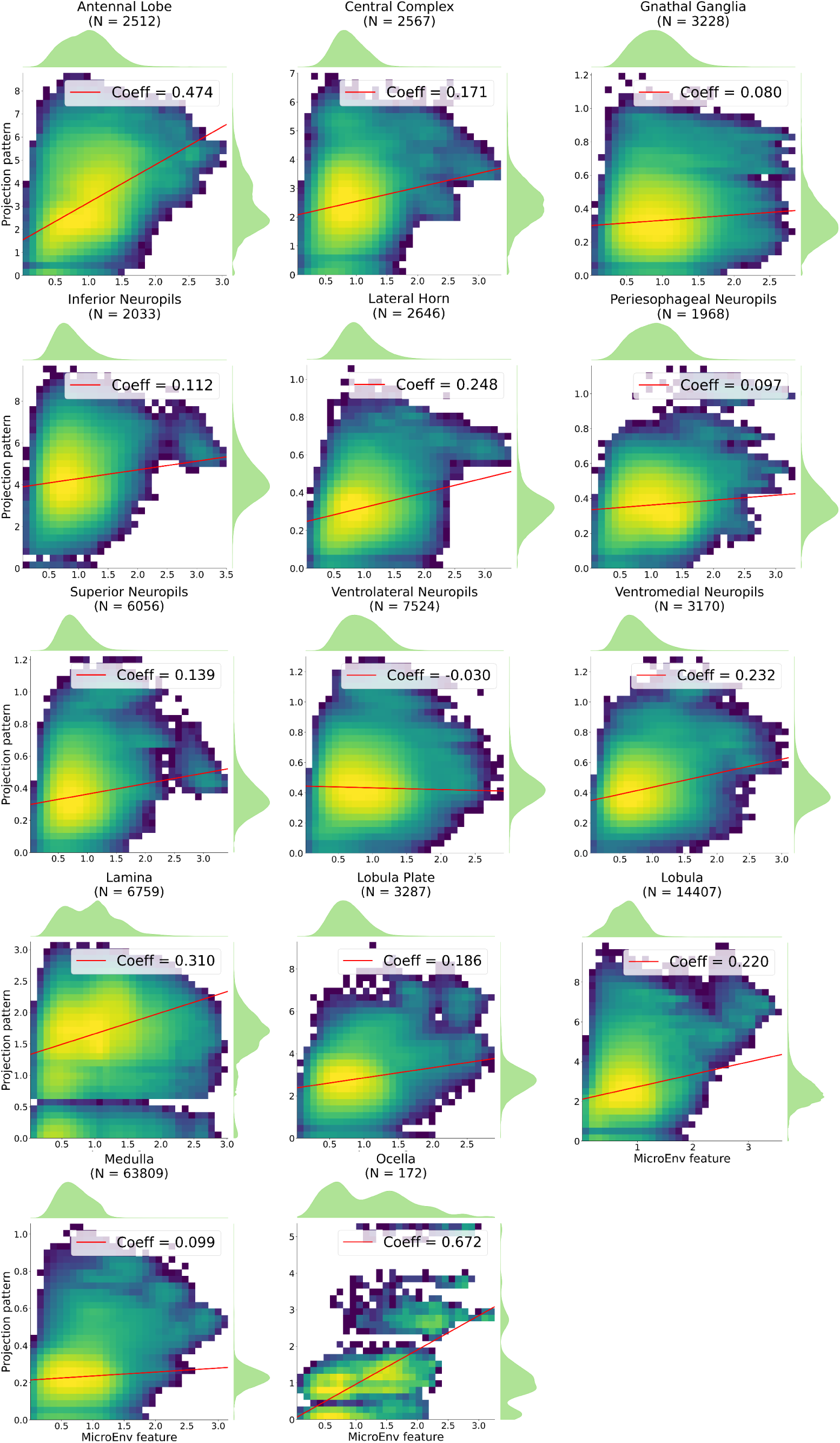
Pearson Correlation Analysis of Pairwise Distances Between Neuronal MicroEnv Features and Axonal Projection Features Across Different Regions in the FAFB Atlas.

**Supplementary Figure 8.**
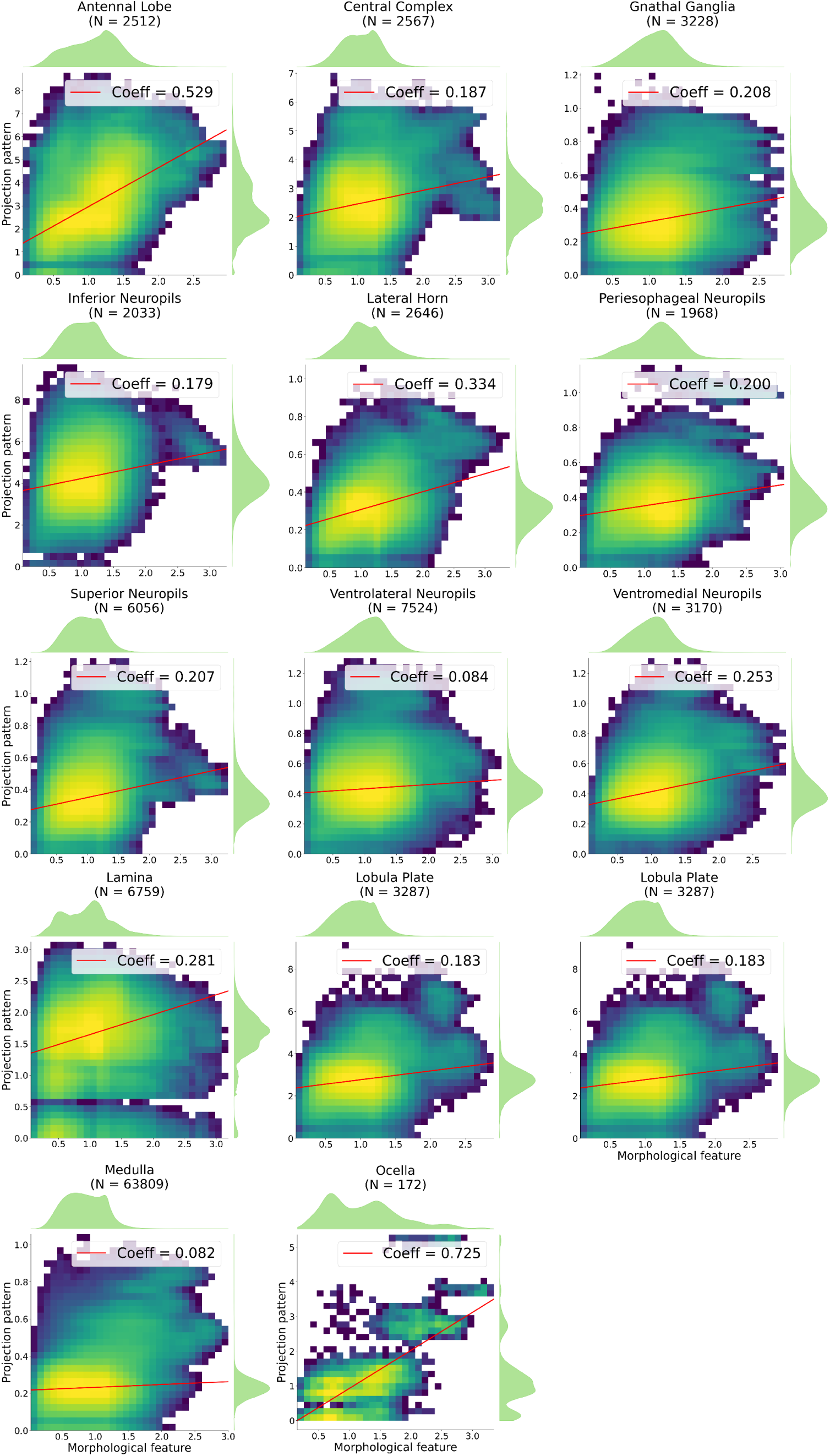
Pearson Correlation Analysis of Pairwise Distances Between Neuronal Morphological Features and Axonal Projection Features Across Different Regions in the FAFB Atlas.

**Supplementary Figure 9.**
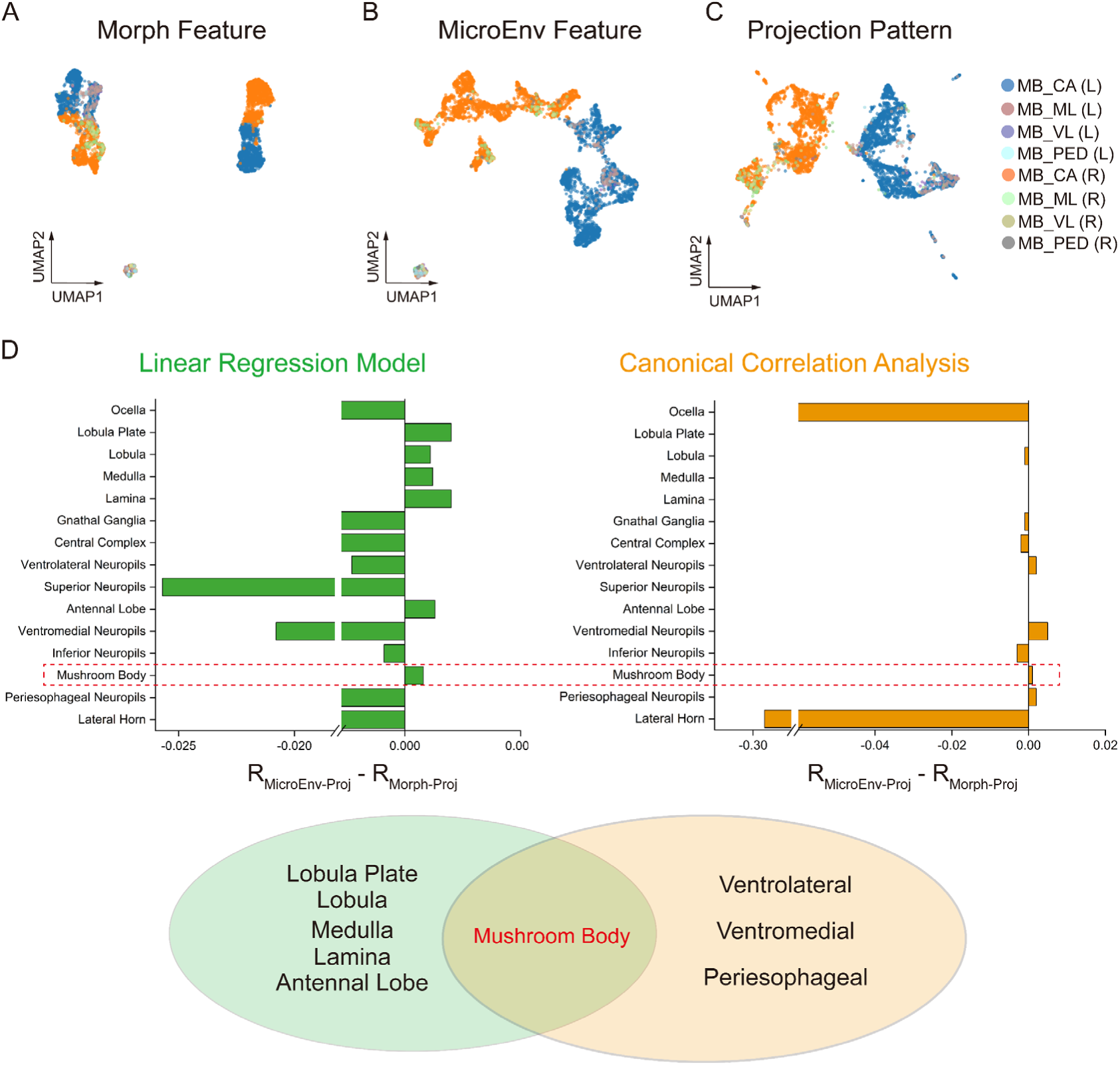
Correlation analysis between MicroE/Morph features and long-range projection features. **A-C.** UMAP two-dimensional visualization of Morph, MicroEnv, and Projection features, with colors representing clusters identified via K-means clustering. **D.** Linear regression analysis of MicroE/Morph features with projection features across different brain regions (left). Canonical correlation analysis (CCA) between MicroE/Morph features and projection features with n_components=3 (right). The x-axis in both panels represents the difference in correlation between MicroEnv and projection features versus Morph and projection features. A Venn diagram (center) illustrates the overlap of brain regions where the correlation difference is greater than 0.

**Supplementary Figure 10.**
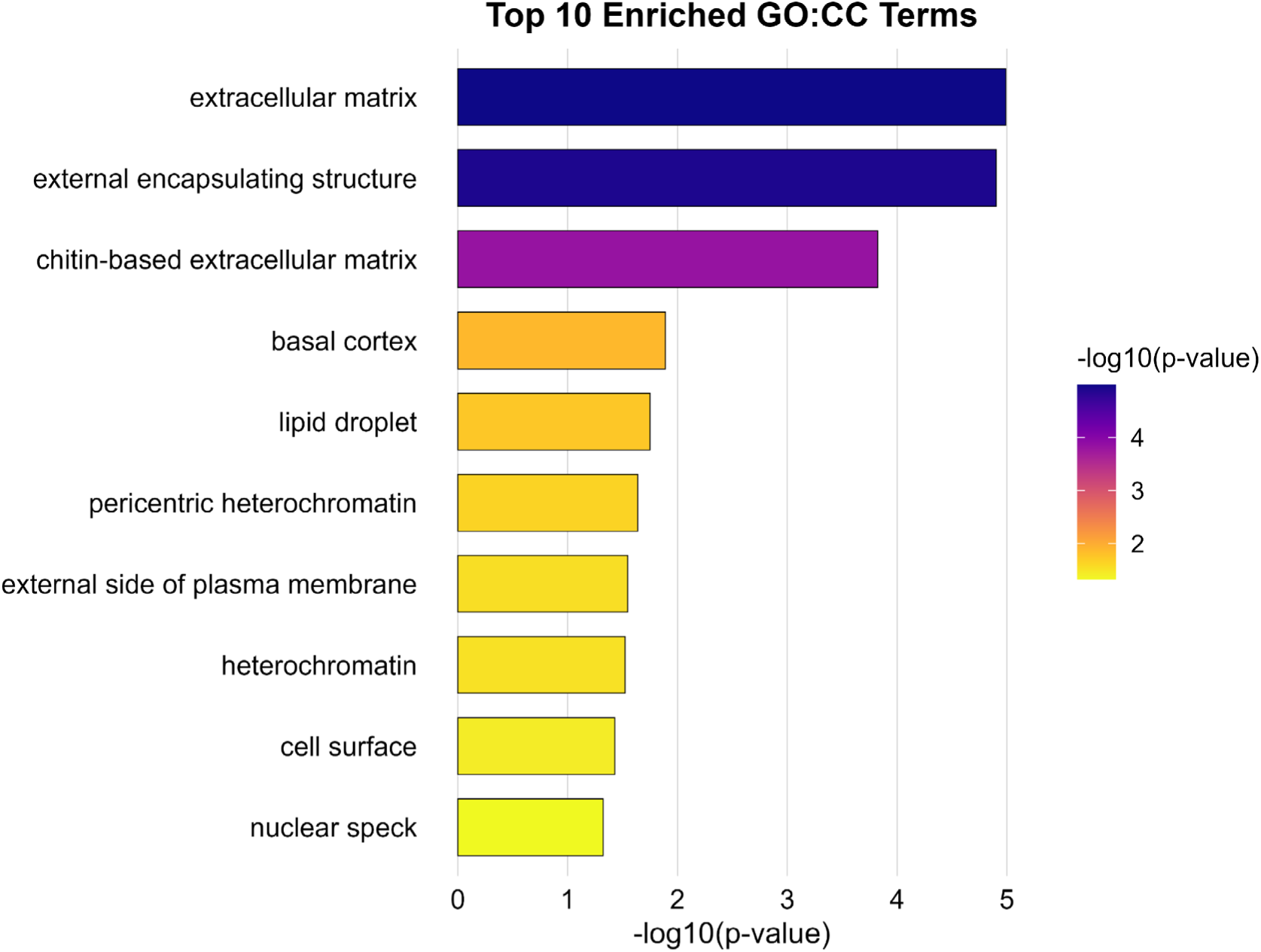
Top 10 Gene Ontology (GO) terms in the Cellular Component (CC) category enriched among the upregulated genes.

**Supplementary Figure 11.**
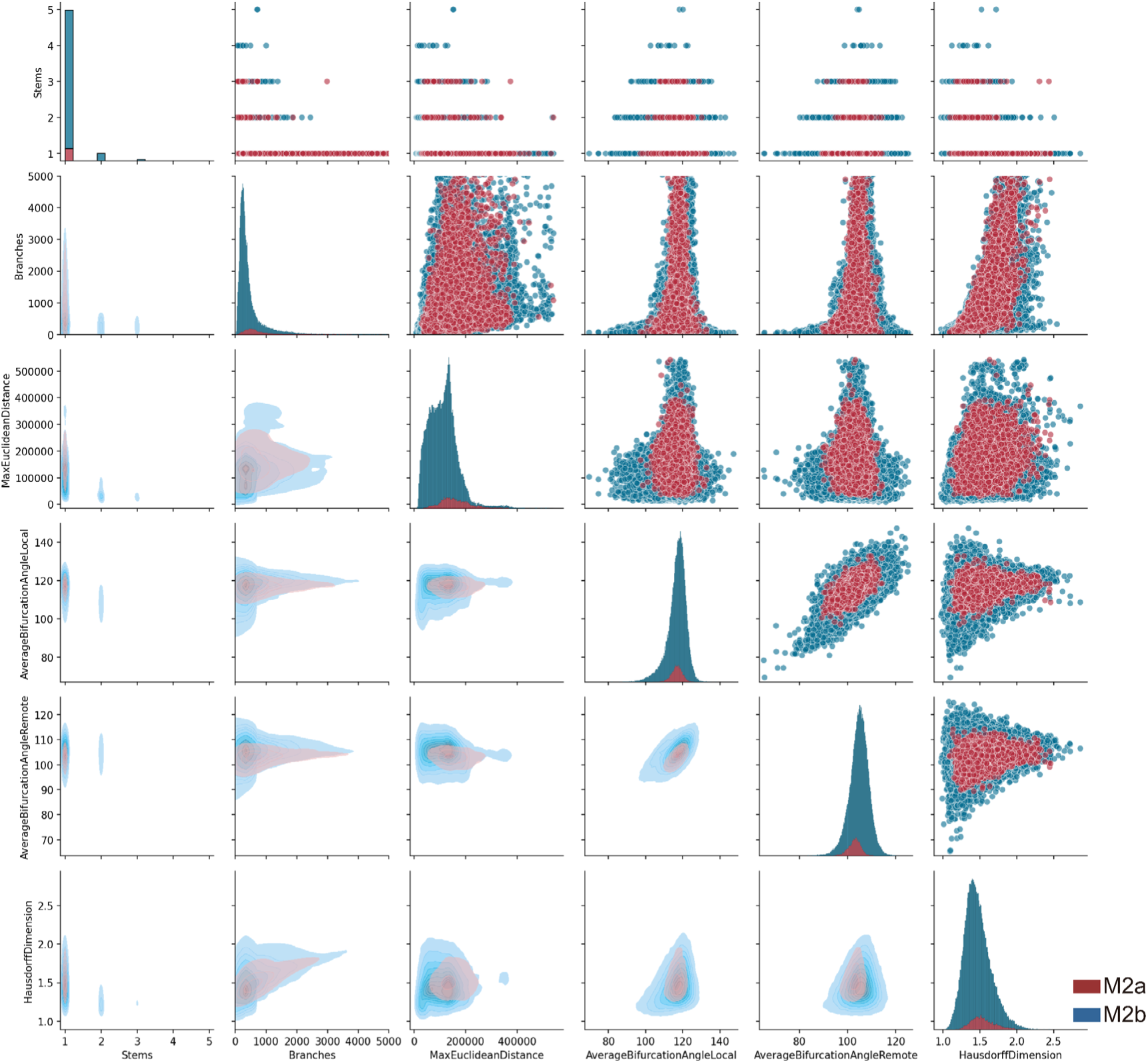
Pair Plot of Morphology Feature Distributions. Pair plot showing the morphology feature distributions for neurons in M2a and M2b, with different colors representing the value distributions of neurons in each module.

**Supplementary Figure 12.**
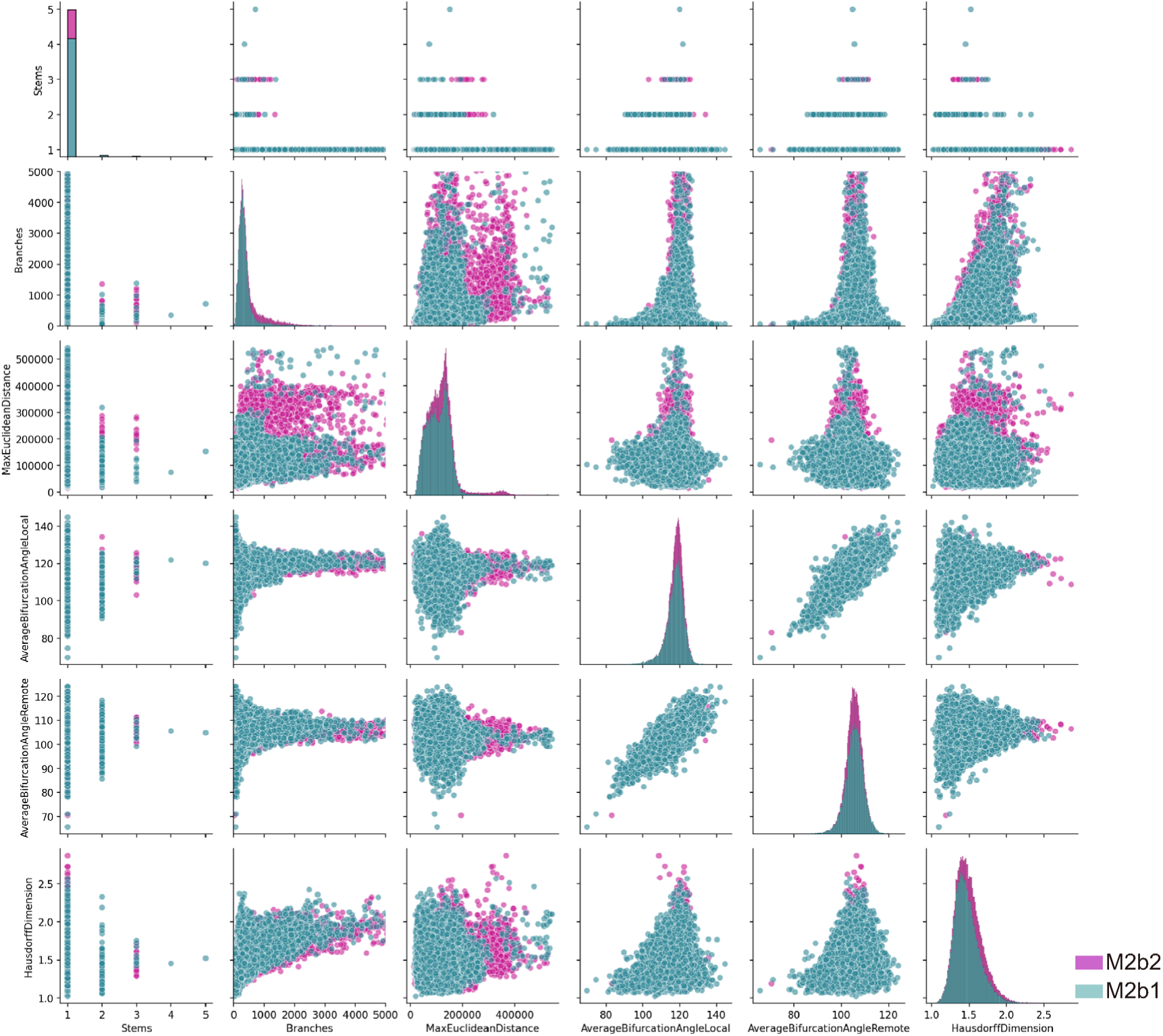
Pair Plot of Morphology Feature Distributions. Pair plot showing the morphology feature distributions for neurons in M2b1 and M2b2, with different colors representing the value distributions of neurons in each module.

**Supplementary Figure 13.**
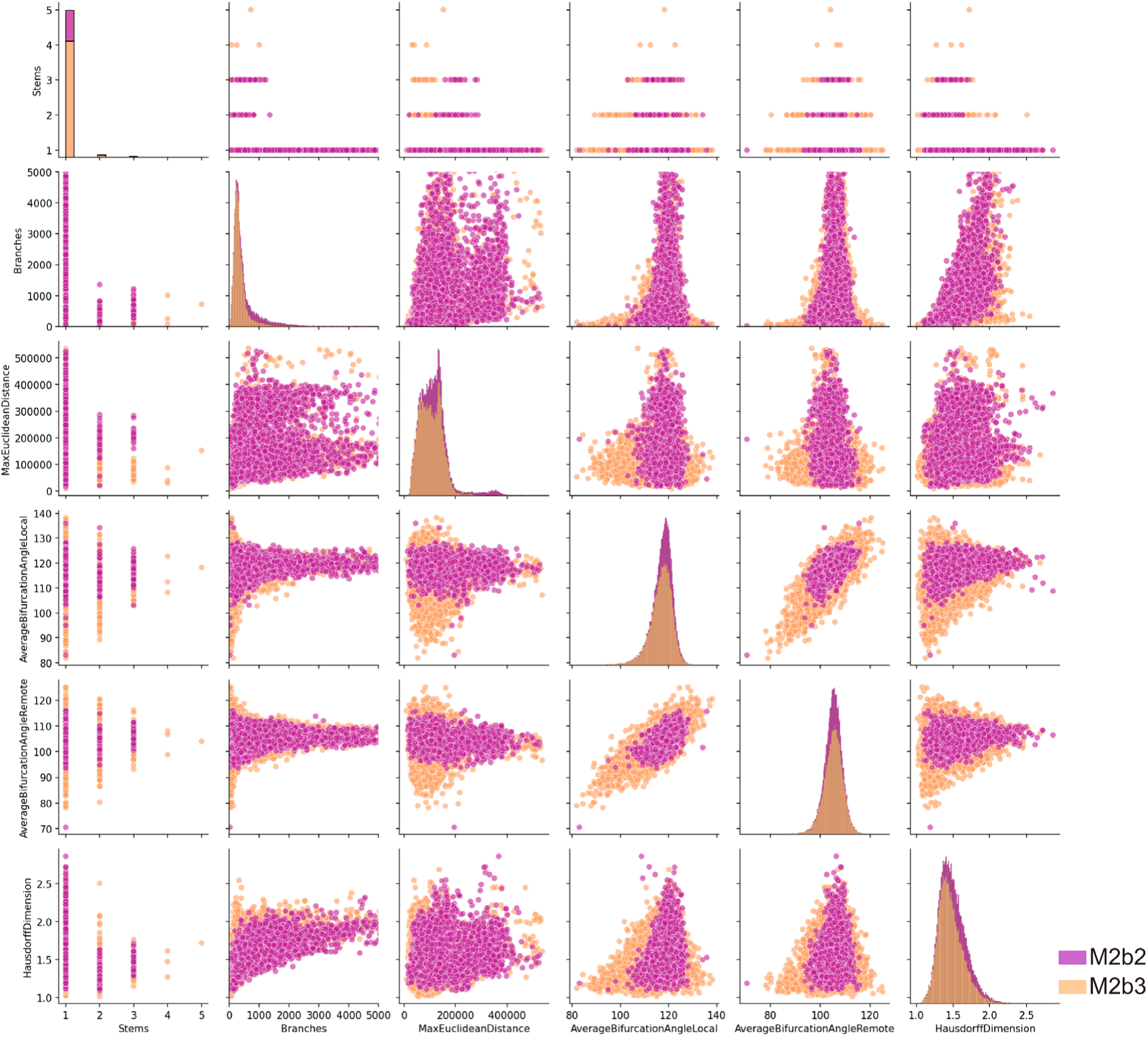
Pair Plot of Morphology Feature Distributions. Pair plot showing the morphology feature distributions for neurons in M2b2 and M2b3, with different colors representing the value distributions of neurons in each module.

**Supplementary Table 1.**
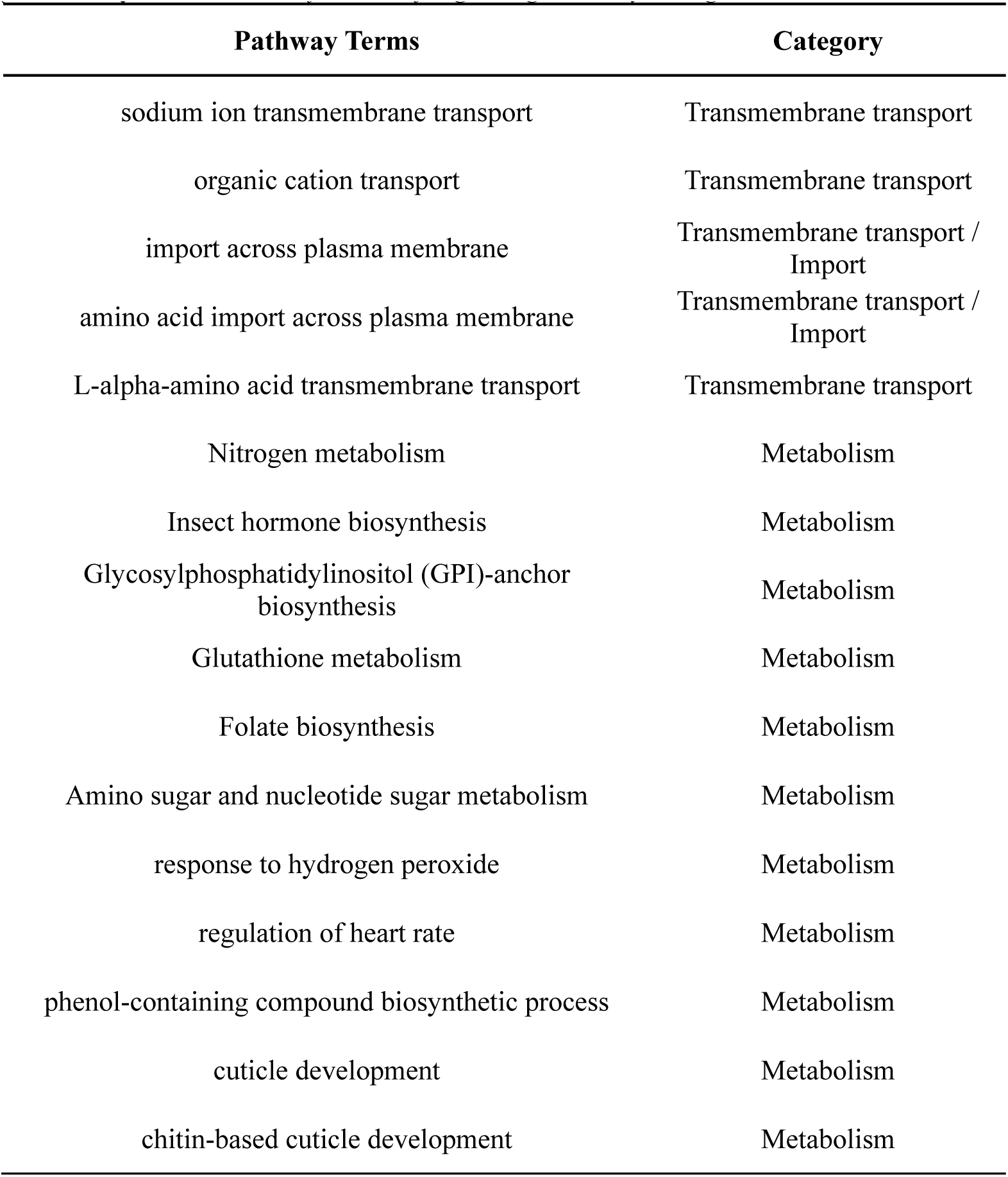
Classification of Signaling Pathway Categories.

